# Mitotic spindle orientation and dynamics are fine-tuned by anisotropic tension via NuMA localisation

**DOI:** 10.1101/2024.10.17.617436

**Authors:** Nawseen Tarannum, Dionn Hargreaves, Dessislava Ilieva, Georgina K. Goddard, Oliver E. Jensen, Sarah Woolner

## Abstract

Cell division orientation, which is key to cell fate and tissue morphogenesis, is influenced by mechanical forces. However, uncoupling whether divisions orient in direct response to force or indirectly via interphase cell shape remains challenging. By applying an external stretch to epithelial tissue and monitoring mitotic spindle dynamics, we show that the spindle orientation protein, NuMA, is recruited to the cell cortex earlier during mitosis in stretched tissue. This stretch-induced recruitment of NuMA coincides with the onset and subsequent amplification of dynamic spindle oscillations. We show by mathematical modelling that increased spindle oscillation indicates increased cortical force generation to orient the spindle. Additionally, we show that knockdown of NuMA reduces spindle oscillations and disrupts division orientation according to stretch and cell shape. Our results indicate that a mechanosensitive re-localisation of NuMA provides a direct response to mechanical force, fine-tuning spindle dynamics and orientation during mitosis and ensuring the accurate alignment of divisions with cell shape and anisotropic tension.

## Introduction

Oriented cell division, dictated by the dynamic positioning of the mitotic spindle, is a key process that contributes to embryonic development. Oriented divisions are important not only for individual cell fates (Cayouette and Raff, 2003; Segalen and Bellaiche, 2009) but also for tissue morphogenesis (Baena-Lopez et al., 2005; Concha and Adams, 1998; Quesada-Hernandez et al., 2010) and tissue homeostasis (Campinho et al., 2013). Therefore, a failure to achieve the correct division alignment leads to abnormalities in morphogenesis and organogenesis (Baena-Lopez et al., 2005; Quesada-Hernandez et al., 2010) and is linked to cancer development in adults (Nakajima et al., 2013; Quyn et al., 2010).

To produce an oriented division, the mitotic spindle is positioned in the cell via dynamic interactions between the spindle’s astral microtubules and core spindle orientation machinery localised at the cell cortex. This machinery canonically consists of a highly conserved ternary complex of Gαi-LGN-NuMA (LGN: Leucine-Glycine-Asparagine-enriched protein; NuMA: nuclear mitotic apparatus protein) (Du and Macara, 2004; Lechler and Fuchs, 2005; Williams et al., 2011) which recruits and anchors dynein at the cortex providing a pulling force to position the spindle (Kotak et al., 2012). The placement of these “cortical force generators” therefore determines the orientation of the spindle and the ultimate division plane. Moreover, cortical force generation also leads to dynamic movements of the mitotic spindle and notably the existence of stereotypical oscillations of the metaphase spindle that coincide with achieving the final division alignment (Grill and Hyman, 2005; Grill et al., 2005; Hargreaves et al., 2024; Larson and Bement, 2017). Mathematical modelling shows that changes to the oscillations of the mitotic spindle can be used as a read out of the factors influencing spindle positioning, such as the density of cortical force generators pulling the spindle into position, or the relative restoring forces generated by growing microtubules (Grill et al., 2005; Hargreaves et al., 2024).

Overlaying the cell-intrinsic spindle orientation machinery is the external environment of the cell, and division orientation has been shown to be highly sensitive to external mechanical force. For example, when tissues are stretched, most divisions orient along the axis of greatest tensile stress (Campinho et al., 2013; Hart et al., 2017; Nestor-Bergmann et al., 2019; Wyatt et al., 2015). However, the molecular mechanism for how forces are sensed to orient divisions remains largely uncharacterised. Forces can be sensed either directly or indirectly – via cell shape changes – to orient divisions. A default mechanism of division orientation, seen in many cell types, is the alignment of the mitotic spindle along the long axis of cell shape, as dictated by Hertwig’s rule (Hertwig, 1884). Since stretch elongates cell shape along the axis of stretch, many studies have reported a cell-shape sensing mechanism for mechanosensitive division orientation (Godard et al., 2020; Lam et al., 2020; Mao et al., 2013; Minc et al., 2011; Nestor-Bergmann et al., 2019; Tang et al., 2018; Thery et al., 2005; Wyatt et al., 2015). However, a growing body of work has also shown that force can be sensed independently of cell shape to orient divisions (Finegan et al., 2019; Fink et al., 2011; Hart et al., 2017; Kelkar et al., 2022; Lisica et al., 2022; Scarpa et al., 2018). A generalised mechanism therefore seems unlikely to capture the robustness of mechanosensitive division orientation. Indeed, a recent study has indicated that both force and cell-shape sensing can act in parallel within the same tissue to position the mitotic spindle (Wang et al., 2017).

For cell-shape sensing, it is thought that cells align divisions to their interphase shape rather than mitotic shape because most cells lose shape anisotropy when they round-up upon mitotic entry (Kunda et al., 2008). In the *Drosophila* pupal notum, the memory of interphase shape is retained during mitosis by the core spindle orientation protein, Mushroom body defect (Mud), the *Drosophila* homologue of NuMA. In this tissue, Mud localises to the tricellular junctions (TCJs), and this localisation pattern is maintained during mitotic rounding providing a blueprint of interphase shape (Bosveld et al., 2016). However, Mud is not enriched at the TCJs in other *Drosophila* tissues (Camuglia et al., 2022; Finegan et al., 2019; Scarpa et al., 2018), raising the question of how divisions are oriented in these tissues. Additionally, Mud has key differences with its vertebrate homologue, NuMA. While Mud is localised to the cortex throughout the cell cycle (Bowman et al., 2006; Izumi et al., 2006; Siller et al., 2006), NuMA has a more dynamic spatiotemporal localisation and is localised to the nucleus during interphase (Kiyomitsu and Boerner, 2021). Therefore, whether Mud-like mechanisms of shape-mediated spindle orientation translate to NuMA in vertebrates is an open question. We have previously shown that LGN, the binding partner of NuMA (Du and Macara, 2004), is concentrated at the tricellular vertices (TCVs; the meeting point of 3 or more cells) throughout mitosis in the *Xenopus laevis* animal cap (Nestor-Bergmann et al., 2019). This TCV localisation of LGN corresponds with the ability of divisions to align with cell shape in this tissue (Nestor-Bergmann et al., 2019). However, whether NuMA localises, similarly to LGN, at the TCVs to orient divisions with cell shape remains unexplored.

Since the mitotic spindle is highly dynamic, a strong possibility is that, following the initial placement along the interphase shape axis, the spindle is reoriented via external cues, such as mechanical force. Indeed, recent studies have reported that the spindle may be dynamically repositioned during metaphase, independent of interphase shape although the mechanism controlling such repositioning remains unclear (Blanchard et al., 2024; Fink et al., 2011; Scarpa et al., 2018). When considering a mechanism that ties spindle orientation with cell shape and external mechanical forces, NuMA emerges as a strong candidate. Indeed, previous studies on fixed keratinocytes have indicated that NuMA could be important for stretch-oriented divisions (Seldin et al., 2013). However, the dynamic role of NuMA in stretch and cell shape-oriented divisions remains uncharacterised especially in the context of complex living tissues.

Here, we investigate the role of NuMA in mechanosensitive spindle orientation using a combination of tissue mechanics, mathematical modelling, and live tissue imaging. We use the *Xenopus laevis* embryonic animal cap, a multilayered vertebrate epithelial tissue that maintains its tissue architecture *ex vivo* and is amenable to mechanical manipulation and imaging (Goddard et al., 2020). We discover that the cortical localisation of NuMA is sensitive to uniaxial stretch with earlier recruitment to the polar cortex during mitosis in stretched tissues. This temporal cortical recruitment is found to coincide with the onset and subsequent amplification of spindle oscillations, which we show by mathematical modelling to indicate an increase in cortical force generation. Crucially, knockdown of NuMA leads to reduced spindle oscillation and reduced accuracy of division alignment with both anisotropic tension and cell shape. Overall, we demonstrate that NuMA specifically links anisotropic tension with division orientation by fine-tuning the dynamics and position of the spindle during mitosis, thus ensuring an accurate alignment of division with cell shape and anisotropic tension.

## Results

### Cortical localisation of NuMA is dynamic and cell cycle dependent

To understand whether NuMA plays a role in mechanosensitive division orientation, we focussed on cortical NuMA since it is the NuMA-dynein complex at the cortex which exerts the pulling forces on astral microtubules to orient the spindle (Kotak et al., 2012). Previous studies have reported that the localisation of NuMA varies throughout the cell cycle (Kiyomitsu and Cheeseman, 2013; Kotak et al., 2013; Seldin et al., 2013; Zheng et al., 2014). Indeed, we were able to visualise this dynamic spatiotemporal localisation by live-imaging GFP-tagged NuMA in the *Xenopus* animal cap, and looking at cell divisions in the superficial layer of this tissue (**Video 1**). During interphase, NuMA accumulates in the nucleus (**Fig. 1A**; **Video 2**). However, when the nuclear envelope breaks down at the onset of mitosis, NuMA is released from the nucleus and localises to the spindle poles and weakly at the cortex (**Fig. 1A**; **Video 2**). The cortical localisation of NuMA becomes more prominent as the cell progresses through mitosis, reaching a peak during anaphase at the cortex closest to the spindle poles. Following anaphase, NuMA is recruited away from the cortex and relocalises to the nuclei of the daughter cells (**Fig. 1A**; **Video 2**).

**Fig. 1.**
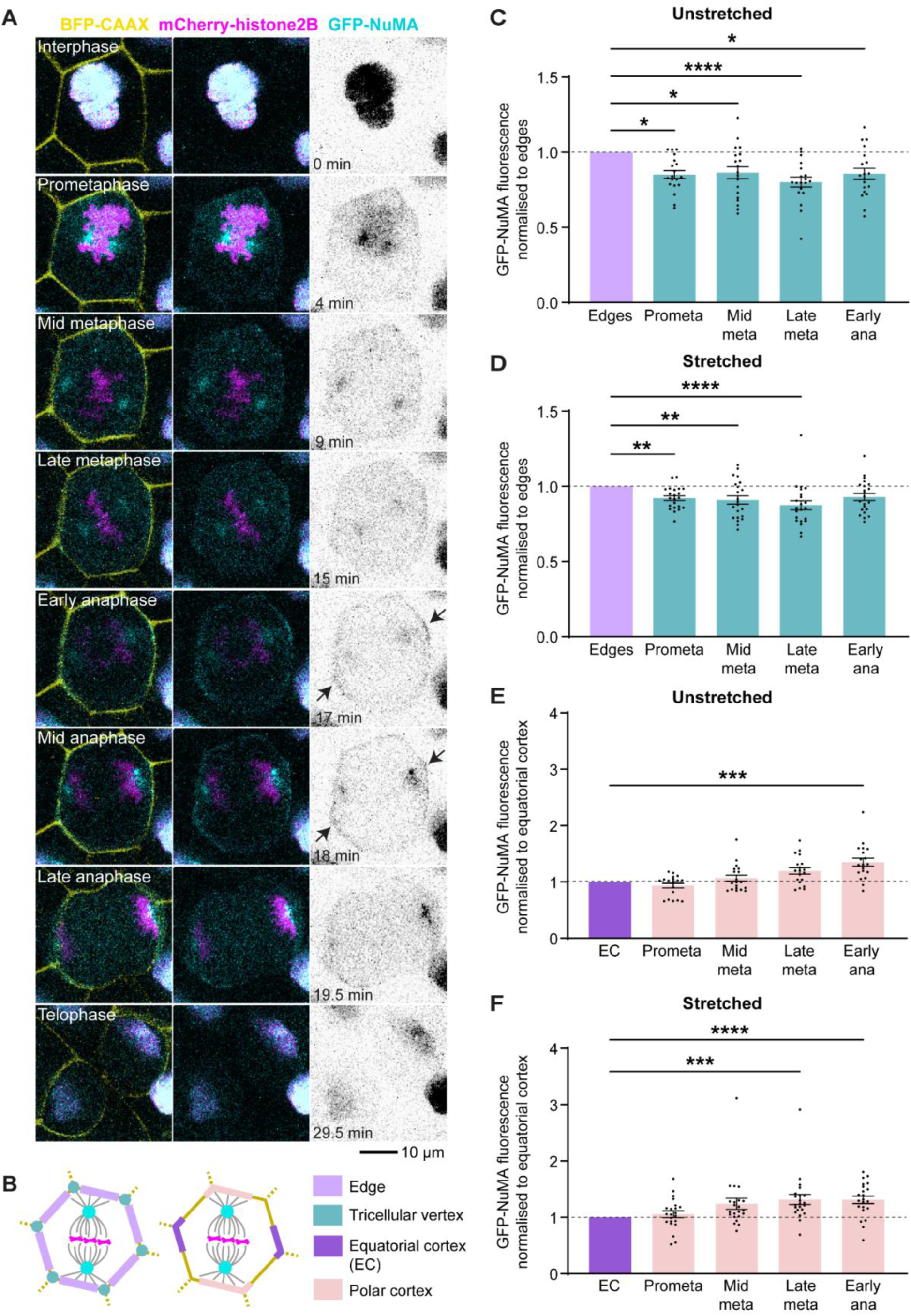
NuMA localises to the cortex earlier in mitosis in tissues under uniaxial stretch. **(A)** A representative cell at interphase and mitosis expressing GFP-NuMA (cyan/inverted grayscale), a membrane marker (BFP-CAAX; yellow), and chromatin marker (mCherry-histone2B; magenta) in an unstretched tissue. Black arrows indicate pronounced cortical localisation of GFP-NuMA. Time is chosen such that 0 minute coincides with the end of interphase. Related to Video 2. **(B)** Fluorescence measurements of GFP-NuMA were taken at the tricellular vertices, cell edges, polar, and equatorial cortex during mitosis. Polar and equatorial cortex were measured using a polygon ROI. **(C, D)** Mean normalised GFP-NuMA fluorescence at tricellular vertices relative to cell edges in unstretched and stretched tissues respectively. Respective non-normalised data are shown in Fig. S1A, B. **(E, F)** Mean normalised GFP-NuMA fluorescence at polar cortex relative to equatorial cortex (polygon ROI) in unstretched and stretched tissues respectively. Respective non-normalised data are shown in Fig. S1C, D. Data in C, D, E, and F were analysed using Friedman test and Dunn’s multiple comparisons test. For all data, n=19 cells, 5 embryos (unstretched); 22 cells, 6 embryos (stretched). Statistical significance: p<0.0001****, p<0.001***, p<0.01**, p<0.05*. All error bars represent mean ± SEM.

### NuMA is not enriched at the tricellular vertices

To quantify the dynamics of GFP-NuMA localisation, we measured the fluorescence of GFP-NuMA at the cortex of each dividing cell at different stages of mitosis. Previously, it was shown that Mud (the *Drosophila* homologue of NuMA) localises to TCJs in the pupal notum tissue (Bosveld et al., 2016). Therefore, we investigated whether, in the *Xenopus* animal cap, NuMA localises similarly to the tricellular vertices (TCVs), where three or more cells meet ( **Fig. 1B**). Moreover, we investigated whether this NuMA localisation was sensitive to mechanical force by measuring GFP fluorescence in both unstretched tissues and in tissues under a uniaxial stretch, applied using an approach described previously (Goddard et al., 2020; Nestor-Bergmann et al., 2019). Interestingly, our data revealed that, in contrast to Mud, NuMA is not significantly enriched at the TCVs relative to the cell edges in either uniaxially stretched or unstretched tissues (**Fig. 1C, D**; non-normalised data in **Fig. S1A**, **B**). In fact, we find that NuMA localisation is significantly reduced at the TCVs compared to the cell edges. Additionally, there was no difference in this localisation pattern between stretched and unstretched tissues (**Fig. 1C, D**; non-normalised data in **Fig. S1A**, **B**).

### Cortical localisation of NuMA is sensitive to tissue stretch

In addition to the TCVs, we measured GFP-NuMA fluorescence at two other regions of the cortex - the polar cortex (closest to the poles) and the equatorial cortex (closest to the metaphase plate) (**Fig. 1B**). We chose these regions to measure as, by eye, we could observe an enhanced cortical localisation of GFP-NuMA at the polar cortex during anaphase (**Fig. 1A, Video 2**). Our quantification indeed revealed that, in unstretched tissues, GFP-NuMA was significantly enriched at the polar cortical regions at early anaphase (**Fig. 1E**; non-normalised data in **Fig. S1C**). Moreover, when a stretch was applied to the tissue, this preferential enrichment of GFP-NuMA at the polar cortex occurred earlier, at late metaphase, and was maintained throughout early anaphase (**Fig. 1F**; non-normalised data in **Fig. S1D**). To prevent selection bias, we used two different ROIs (polygon and oval) to measure the fluorescence of cortical NuMA. Importantly, the cortical localisation trend of NuMA observed between unstretched and stretched tissues was similar irrespective of the type of ROI used for fluorescence measurements (**Fig. S1E**, **G**, **H**). Additionally, we did not observe any change to the total fluorescence of GFP-NuMA at the cortex between stretched and unstretched tissues (**Fig. S1E**, **F**). This suggests that rather than an increase in overall cortical NuMA, there is a redistribution of cortical NuMA between the equatorial and polar cortex in the presence of tissue stretch. Together, these data demonstrate that rather than being enriched at the TCVs, NuMA is localised preferentially to the polar cortex. Uniaxial stretch, in turn, facilitates an earlier recruitment of NuMA at the polar cortex during mitosis.

### Spindle oscillations are amplified by mitotic progression and by stretch

NuMA is known to function at the cell cortex as part of a complex with dynein which generates pulling forces on astral microtubules to position the spindle (Kiyomitsu and Cheeseman, 2013; Okumura et al., 2018). Mathematical modelling by our group and others has shown that the combination of the attachment and detachment of these “cortical force generators” and the antagonistic centring forces of spindle microtubules results in dynamic movements of the spindle pole (Grill et al., 2005; Hargreaves et al., 2024). By examining these dynamic spindle movements, we aimed to determine whether the temporal recruitment of NuMA to the cell cortex aligns with observed spindle movements, thereby implicating NuMA’s cortical localisation as a leading factor in spindle positioning under tension.

Coordinated dynamic movements of both spindle poles can result in rotational movements of the mitotic spindle. These movements can be split into two categories: overall spindle rotation from metaphase onset until the end of anaphase; and the finer movements which overlay the larger-scale movement. These finer movements may be oscillatory (**Fig. 2A; Video 3**) or non-oscillatory (**Video 4**). Both rotational movements have been observed in the animal cap at this stage (Hargreaves et al., 2024; Larson and Bement, 2017). Crucially, mathematical modelling of spindle pole dynamics has shown that the oscillatory movement of the spindle is dependent on the number of cortical force generators (*N*) at the cortex, with small increases in *N* promoting a switch between non-oscillatory and oscillatory movement (**Fig. 2B**) (Grill et al., 2005; Hargreaves et al., 2024). Stochastic binding and unbinding of force generators promotes the onset of noise-induced oscillations outside of the stability boundary (purple shaded region to the left of the stability line, **Fig. 2B**), making oscillations viable at smaller values of *N* (Hargreaves et al., 2024). The same model can be used to predict the relationship between oscillation amplitude and *N* (**Fig. 2C**), which is also extended at the stability boundary by the presence of noise. For a constant binding rate between cortical force generators and microtubules, a positive relationship between *N* and the amplitude of oscillation is predicted (**Fig. 2C**) (Hargreaves et al., 2024). The period of oscillation around the stability boundary is also shown to vary with *N* (**Fig. 2D**). Our observation that NuMA is recruited to the polar cortex earlier in stretched tissues (**Fig. 1E, F**), would be predicted to correspond to an increase in *N* earlier in metaphase. We sought to determine if any related changes in spindle movement could be identified that might indicate a change in cortical force generation when the tissue is stretched.

**Fig. 2.**
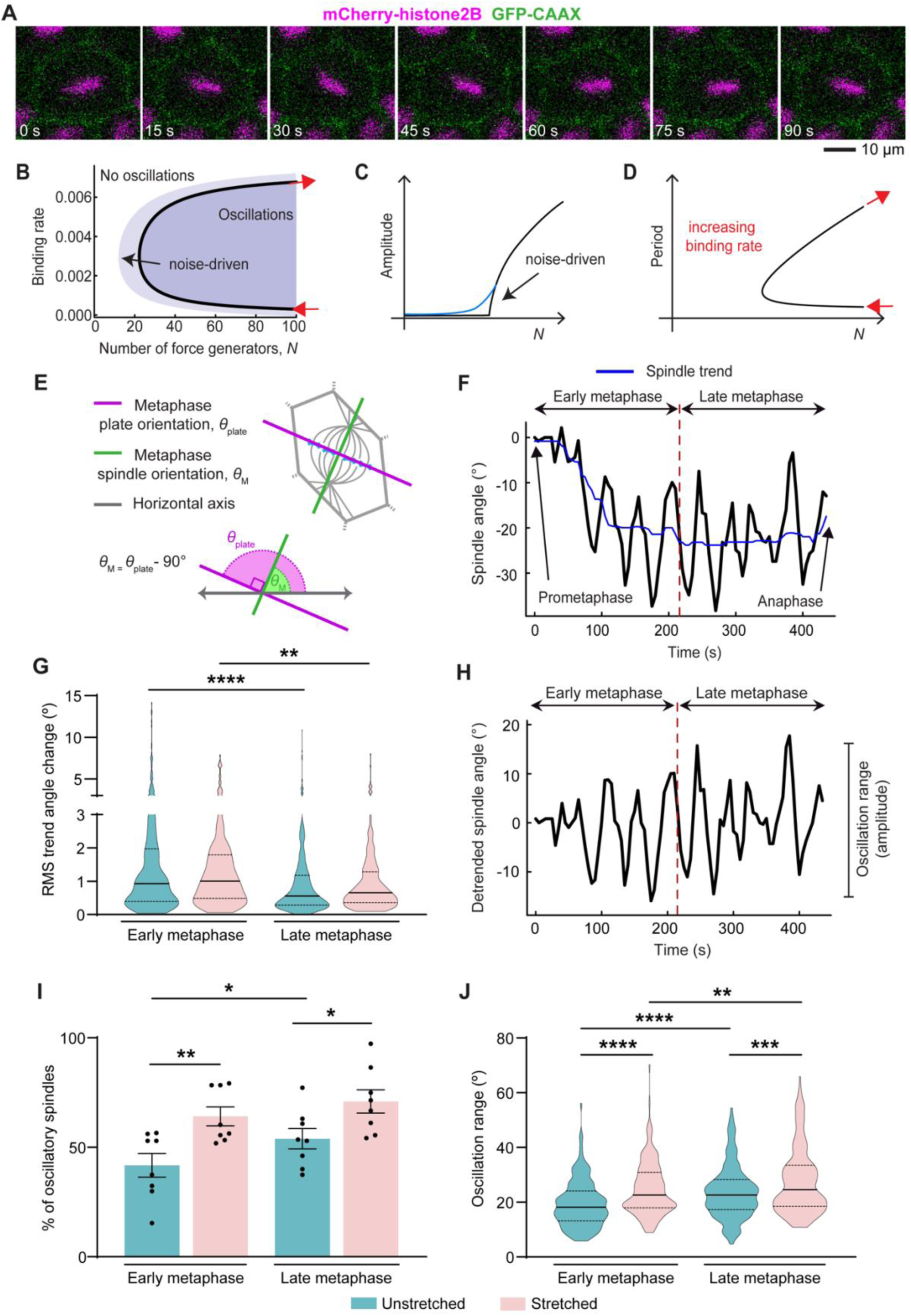
The mitotic spindle oscillates in a stretch-dependent manner. **(A)** A representative cell at metaphase in a stretched tissue expressing a membrane marker (GFP-CAAX; green), chromatin marker (mCherry-histone2B; magenta), and containing control morpholino. Time is chosen such that 0 second coincides with the start of metaphase plate oscillation. One whole period of oscillation is shown between *t*=0 s and *t*=60 s. Related to Video 3. **(B)** Mathematically derived stability boundary (black curve), showing the boundary between oscillatory (blue shaded region) and non-oscillatory spindle pole dynamics. The binding rate between force generators (NuMA/dynein complex) and microtubules is non-dimensional. *N* represents the number of available force generators. The crossing between non-oscillatory and oscillatory solutions is extended in the presence of noise, where noise-induced oscillations may arise. Red arrows identify the path followed along the stability boundary in C (Hargreaves et al., 2024). **(C)** Amplitude of oscillation increases with *N* for a constant binding rate. The amplitude increases from 0 as *N* crosses the stability boundary indicated in A. This sharp transition is smoothed in the presence of noise due to noise-induced oscillations (blue line) (Hargreaves et al., 2024). **(D)** Relationship between the period of oscillation and *N,* along the stability boundary identified in A. The direction of increasing binding rate along the stability boundary is indicated in red (Hargreaves et al., 2024). **(E)** Schematic showing the measured orientation of the metaphase plate (*θ*_plate_) and the corresponding metaphase spindle orientation (*θ*_M_) lying perpendicular to one another. **(F)** Track of a representative spindle undergoing rotational movements from prometaphase until the start of anaphase. Red dashed line corresponds to the mid metaphase timepoint. The overall spindle movement (blue line) is determined using a moving median with a window length Δ*t*=125 s. **(G)** Root mean square (RMS) of the change in the trend angle between successive timepoints in unstretched and stretched tissues in early and late metaphase. Data analysed using a mixed-effects model and Tukey’s multiple comparisons test. **(H)** The spindle track from C with the trend (blue) removed. The measured oscillation range is defined by the upper and lower bounds of the signal omitting outliers. **(I)** Percentage of dividing cells with oscillatory spindles in stretched and unstretched tissues, in early and late metaphase. Data were analysed using two-way ANOVA and Fisher’s least significant difference test. Error bars represent mean ± SEM. For D and F, n=342 cells, 8 embryos (unstretched); 237 cells, 8 embryos (stretched). **(J)** Range of oscillation for oscillatory spindles in stretched and unstretched tissues, in early and late metaphase. Data were analysed using a mixed-effects model and Fisher’s least significant difference test. n=152 cells, 8 embryos; 193 cells, 8 embryos (unstretched, early and late metaphase respectively); 156 cells, 8 embryos; 174 cells, 8 embryos (stretched, early and late metaphase respectively). For D and G, solid and dashed lines represent median and quartiles respectively. Statistical significance: p<0.0001****, p<0.001***, p<0.01**, p<0.05*.

To track spindle movements, we used the position and orientation of the metaphase plate (**Fig. 2A**), which we found to be an accurate proxy for spindle movement (**Fig. S2A-F**). The angle of the metaphase plate was shown to lie perpendicular to the angle of the metaphase spindle (89.7 ± 0.3°, mean ± SEM; **Fig. S2E**) with highly correlated inter-frame angle displacements (**Fig. S2D**, **F**). We measured the rotational movement of mitotic spindles from metaphase onset until anaphase by following the metaphase plate angle in time (**Fig. 2E, F**; **Fig. S2G**, **H, J**). As expected, we typically observed an overall spindle rotation or trend (**Fig. 2F** blue line), superimposed with finer movements. To determine how spindle dynamics change over the course of mitosis, we compared movements between early metaphase (defined as the first half of metaphase) and late metaphase (the second half of metaphase) (**Fig. 2F**). We first analysed the spindle trend (**Fig. 2F** blue line) to determine whether the overall spindle rotation towards the ultimate division axis was impacted by the application of stretch. By calculating the root mean square (RMS) of the angular displacement between sequential timepoints, we saw biphasic behaviour whereby the spindle moves most in early metaphase, followed by a reduction in overall rotation in late metaphase (**Fig. 2G**). Notably, this spindle rotation trend was unaffected by the application of stretch. This suggests that the application of stretch does not affect the overall rotation of the spindle toward its final orientation.

We next focussed on the smaller movements overlaying the overall rotation by removing the trend line from the spindle angle measurements (**Fig. 2H**). Spindles were determined to be oscillatory (**Fig. S2H; Video 3**) or non-oscillatory (**Fig. S2J; Video 4**) using an autocorrelation function (**Fig. S2I, K**). The presence of significant peaks in correlated movements at increasing time-lags was used as an indicator for spindle oscillations (**Fig. S2I, K**). In unstretched tissues we observed an increase in the percentage of spindles undergoing oscillatory movements as mitosis progresses, with significantly more spindles undergoing oscillatory movements in late compared to early metaphase (**Fig. 2I**). Moreover, in stretched tissues we saw a significantly larger fraction of spindles undergoing oscillatory movements in both early and late metaphase compared to unstretched tissues. Notably, in stretched tissues, the percentage of oscillatory spindles in early metaphase was very similar to that seen in unstretched tissues in late metaphase (64 ± 4% and 54 ± 5%, respectively, mean ± SEM; **Fig. 2I**), indicating that the increase in oscillatory spindles normally seen as mitosis progresses is accelerated by stretch.

To further investigate the dynamics of spindle oscillations, we defined an oscillation range by taking the upper and lower bounds of the signal, omitting outliers. The range spanned by the remaining signal provides a measure related to the oscillation amplitude (**Fig. 2H**, oscillation range indicator). As mathematical modelling suggests that increasing *N* would result in an increase in the amplitude of a spindle pole oscillation (**Fig. 2C**), we utilised this measurement as an indicator of whether changes in the spindle movements could be a result of the temporal accumulation of NuMA. We observed that the oscillation range (amplitude) significantly increased between early and late metaphase in both stretched and unstretched tissues (**Fig. 2J**). Moreover, the oscillation range was observed to be significantly increased in stretched tissues compared to unstretched tissues in both early and late metaphase (**Fig. 2J**). In addition, just as we saw with the percentage of oscillatory spindles, the oscillation range seen in stretched tissues at early metaphase was very similar to the range seen in late metaphase in unstretched tissues (24.8 ± 0.8° and 23.7 ± 0.7°, respectively (mean ± SEM); **Fig. 2J**).

The mathematical model highlights that the period of oscillation around the neutral curve has a non-linear relationship with *N* (**Fig. 2D**). Experimentally, we observed that the application of stretch increased the period of oscillation from 66 ± 2 s to 72 ± 2 s (mean ± SEM) in late metaphase (**Fig. S2L**). In early metaphase we observed no significant difference in period between stretched and unstretched conditions. This general trend is congruent with increasing *N* leading to an increased period of oscillation (upper branch of **Fig. 2D**). We highlight that the measurement of the period of oscillation is sensitive to noise, and less accurate for shorter time series. As the time series length was limited by the length of metaphase, we use the oscillation range as a more comparable and measurable change to the spindle oscillations.

Overall, our analysis of spindle rotational movement indicates that stretching accelerates and accentuates the changes in spindle oscillations normally seen as mitosis progresses. This acceleration in the oscillatory dynamics of spindles in stretched tissues mirrors the earlier recruitment of NuMA to the polar cortex that we see following stretch and indicates that the cortical forces acting on the spindle increase when the tissue is stretched.

### Knockdown of NuMA affects embryonic development and spindle bipolarity

We next sought to functionally test the role of NuMA in mechanosensitive spindle orientation, by knocking down the expression of endogenous NuMA in early *Xenopus* embryos. Using translation blocking morpholinos (MOs), we achieved a 46% knockdown of NuMA expression at the early gastrula stage (**Fig. 3A**). We then characterised the effects of NuMA knockdown on embryo development. Despite the partial knockdown of NuMA expression via MOs, we observed markedly increased lethality at the end of gastrulation in embryos injected with NuMA MO compared to uninjected embryos and embryos injected with control MO (**Fig. S3A, B**). Therefore, NuMA is an essential gene during *Xenopus* development similar to that in mice (Silk et al., 2009) and in humans (based on data from genetic screens) (Blomen et al., 2015; Georgi et al., 2013; Hart et al., 2015; Wang et al., 2015). Importantly, embryos injected with NuMA MO were healthy at the early gastrula stage (9 h timepoint in **Fig. S3A**) at which the animal cap tissue was dissected for stretch experiments. Since NuMA is important for spindle structure (Silk et al., 2009), we tested the effect of NuMA knockdown on spindle morphology in fixed early gastrula embryos. In line with previous studies (Ma et al., 2022; Magescas et al., 2017), we observed an increased incidence of multipolar spindle formation upon knockdown of NuMA (**Fig. S3C, D**). Importantly, most spindles had a normal structure despite the knockdown (**Fig. S3D**). This allowed us to analyse normal spindles within the knockdown background in subsequent live imaging experiments.

**Fig. 3.**
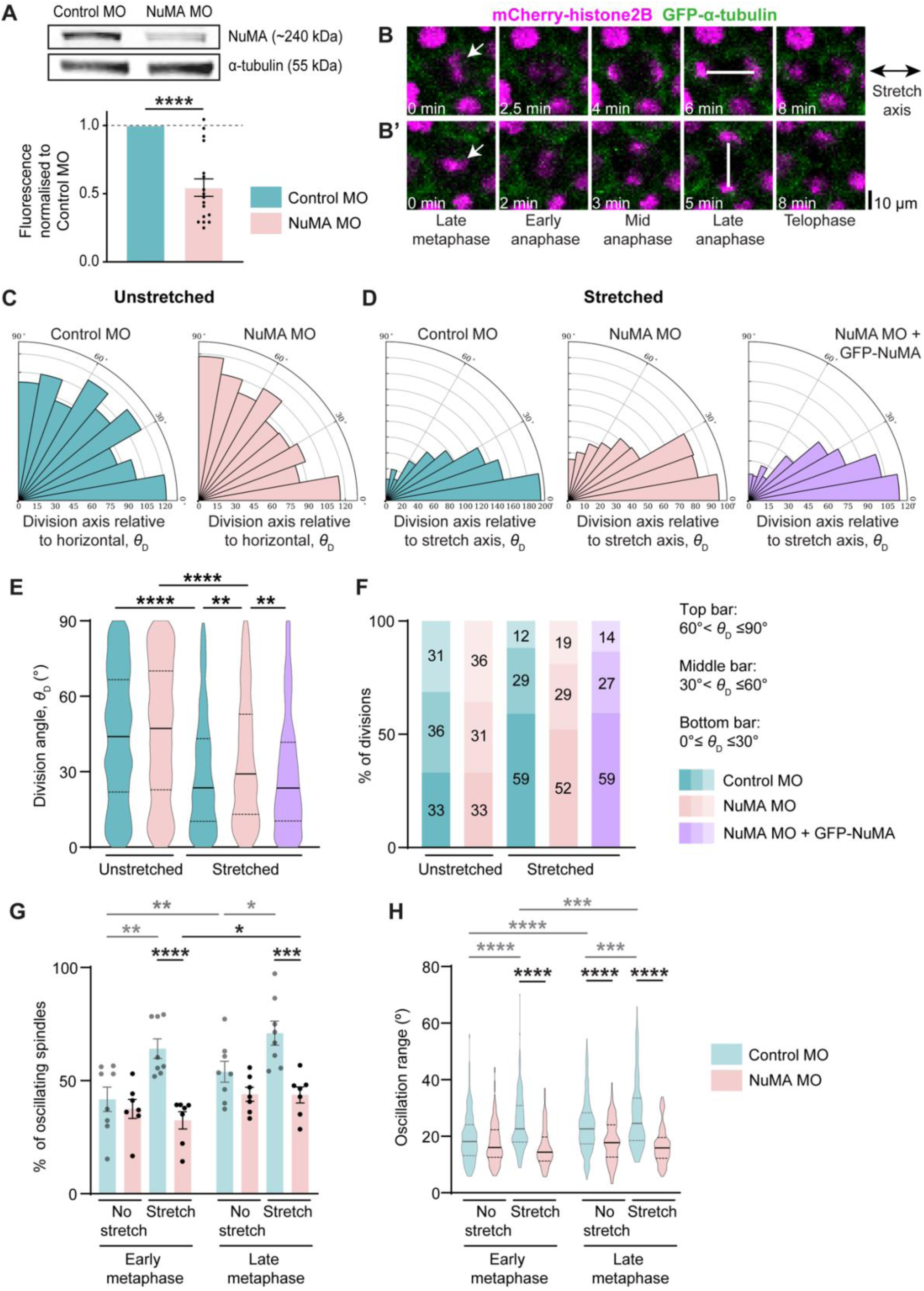
Knockdown of NuMA disrupts division orientation along the axis of stretch and spindle oscillation. **(A)** Endogenous *Xenopus* NuMA expression following knockdown of NuMA. Data analysed using Kolmogorov-Smirnov test. n=17 independent repeats per condition from 170 embryos. **(B)** A representative cell dividing parallel to the axis of stretch (double sided black arrow) or **(B’)** perpendicular to the stretch axis. The cells are expressing a microtubule marker (GFP-α-tubulin; green), chromatin marker (mCherry-histone2B; magenta) and contain control MO. White bars indicate the orientation of division. White arrows indicate the position of the metaphase plate at the end of metaphase. Time is chosen such that 0 minute coincides with late metaphase. The images for B and B’ are related to Video 6 and 7 respectively. **(C, D)** Division angles relative to the horizontal axis (the stretch axis in stretched tissues) in control MO, NuMA MO, and rescue tissues, with the number of divisions in 10° bins. **(E)** Distribution of division angles in the unstretched and stretched control MO, NuMA MO, and rescue tissues. Data were analysed using Kruskal-Wallis test and Dunn’s multiple comparisons test. n=922 cells, 9 embryos; 887 cells, 13 embryos (unstretched, control MO and NuMA MO respectively); 810 cells, 9 embryos; 502 cells, 11 embryos (stretched, control MO and NuMA MO respectively); 495 cells, 11 embryos (rescue). **(F)** Percentage of divisions from each distribution in E binned into three partitions relative to the horizontal axis (the stretch axis in stretched tissues). **(G)** Percentage of dividing cells with oscillatory spindles in stretched and unstretched tissues injected with control MO and NuMA MO, in early and late metaphase. Data were analysed using two-way ANOVA and Šidák’s multiple comparisons test. Control MO data and comparisons as shown previously, shaded grey for clarity. n=342 cells, 8 embryos; 213 cells, 7 embryos (unstretched, control MO and NuMA MO respectively); 237 cells, 8 embryos; 201 cells, 7 embryos (stretched, control MO and NuMA MO respectively). For A and G, error bars represent mean ± SEM. **(H)** Range of oscillation for oscillatory spindles in stretched and unstretched tissues injected with control MO and NuMA MO, in early and late metaphase. Data were analysed using a mixed-effects model and Tukey’s multiple comparisons test. Control MO data and comparisons as shown previously, shaded grey. n=152 cells, 8 embryos; 193 cells, 8 embryos (unstretched control MO, early and late metaphase respectively); 156 cells, 8 embryos; 174 cells, 8 embryos (stretched control MO, early and late metaphase respectively); 87 cells, 7 embryos; 96 cells, 7 embryos (unstretched NuMA MO, early and late metaphase respectively); 67 cells, 7 embryos; 89 cells, 7 embryos (stretched NuMA MO, early and late metaphase respectively). Statistical significance: p<0.0001****, p<0.001***, p<0.01**, p<0.05*. For E and H, solid and dashed lines represent median and quartiles respectively.

### NuMA is required for accurate division orientation with stretch

Previously, NuMA was shown to be important for stretch-dependent spindle orientation in fixed cells (Seldin et al., 2013). Therefore, we tested the extent to which knockdown of NuMA affects stretch-sensitive divisions in live cells within a tissue environment. We did not observe any significant effect of NuMA knockdown on cell division rate (**Fig. S3E**). Therefore, we focussed on how division orientation may be affected by knockdown of NuMA. To do this, we measured the angle between the spindle axis at late anaphase, when the final division orientation is set (Woolner et al., 2008; Woolner and Papalopulu, 2012), and the horizontal axis (the stretch axis in stretched tissues). Within the tissue, cells can divide at a variety of angles relative to the horizontal/stretch axis (**Video 5**). These division angles can range from 0° (**Fig. 3B; Video 6**) to 90° (**Fig. 3B’; Video 7**) relative to the horizontal/stretch axis. In unstretched tissues, there was no significant difference between the distributions of control and NuMA knockdown tissues (**Fig. 3C, E**). In stretched control MO tissues, the majority of divisions oriented with the axis of stretch (within 30°) (**Fig. 3D**), corroborating previous studies showing that application of an external uniaxial stretch orients divisions (Hart et al., 2017; Nestor-Bergmann et al., 2019; Wyatt et al., 2015). This was also the case in stretched NuMA knockdown tissues (**Fig. 3D**). Importantly, however, there was a significant difference between the distribution of division orientation and the stretch axis in the stretched control and NuMA knockdown tissues, with NuMA knockdown tissues showing a small but significant loss of alignment (**Fig. 3D, E**).

To further probe the nuances of each distribution, we partitioned the division angles into three 30° bins relative to the horizontal/stretch axis (**Fig. 3F**). In the unstretched tissues, divisions were uniformly distributed across each bin. In the stretched tissues, most divisions were concentrated within 30° of the stretch-axis. However, while 59% of the divisions oriented within 30° of the stretch-axis in the control tissues, this decreased to 52% in the NuMA knockdown tissues. Conversely, while 12% of divisions oriented at angles greater than 60° in control tissues, this increased to 19% in NuMA knockdown tissues (**Fig. 3F**).

Importantly, this division misalignment with the stretch axis in NuMA knockdown tissues was rescued by expressing GFP-NuMA over the endogenous knockdown background (**Fig. 3D, E, F**). The dose of exogenous GFP-NuMA was carefully titrated to match the level of NuMA expressed endogenously (**Fig. S3F**). Overall, these data show that NuMA is important for mechanosensitive cell division orientation with NuMA knockdown leading to the misalignment of divisions with the stretch axis.

### NuMA is required for stretch-induced spindle oscillation amplification

Given the identification of an accentuation of spindle oscillations under stretch, coinciding temporally with increased GFP-NuMA localisation to the polar cortex, we sought to determine whether NuMA knockdown would impact spindle oscillations accordingly. Indeed, the overall percentage of spindles showing an oscillatory phenotype was significantly decreased upon NuMA knockdown in both early and late metaphase (**Fig. 3G**). The percentage of oscillatory spindles in stretched tissues is almost halved by the partial knockdown of NuMA (64 ± 4% control vs 32 ± 4% NuMA knockdown in early metaphase and 71 ± 5% control vs 44 ± 4% NuMA knockdown in late metaphase; mean ± SEM). Additionally, the percentage of spindles showing an oscillatory phenotype upon NuMA knockdown across both stretched and unstretched conditions remained similar to unstretched control levels (**Fig. 3G**).

Within the population of spindles which did oscillate, the range (amplitude) of the oscillation was also seen to significantly decrease in NuMA knockdown compared to control, upon the application of a tissue stretch in both early and late metaphase (**Fig. 3H**), congruent with an overall reduction of *N* (**Fig. S2B**). Just as with the percentage of spindles oscillating, knockdown of NuMA kept oscillation amplitude in stretched tissue at a level similar to unstretched. Overall, these data show that a partial NuMA knockdown is sufficient to induce significant differences in the dynamic rotation of the mitotic spindle under stretch. In particular, with NuMA knockdown, we observe a dampening down of the normal increases in oscillation (percentage and amplitude) seen when a control tissue is stretched. These results indicate that the increase in spindle oscillation and inferred increase in cortical force generation seen upon tissue stretch (**Fig. 2; Fig. S2**) are dependent upon the presence on NuMA.

### NuMA is required to fine-tune division alignment with cell shape under anisotropic stretch

Previous studies have shown that application of a uniaxial stretch changes the shape of cells, with more elongated cells in stretched tissues and an alignment of the long axis of cell shape with the stretch axis. Since cells are known to preferentially align their divisions with the interphase long axis of shape, the introduction of cell shape anisotropy upon stretch has been suggested to drive alignment of division with tensile force in many tissues (Nestor-Bergmann et al., 2019; Wyatt et al., 2015). We therefore hypothesised that the loss of division alignment with the stretch axis seen upon NuMA knockdown could result from reduced alignment with cell shape more generally. To test this, we explored the effect of NuMA knockdown on division alignment with cell shape.

As expected, we observed an increase in cell elongation upon application of a uniaxial stretch in both control and NuMA knockdown tissues (**Fig. 4A, B**). We determined the principal axis of shape (*θ*_V_) at interphase just before nuclear envelope breakdown, based on the position of the TCVs (Nestor-Bergmann et al., 2019). The orientation of cell division relative to cell shape was then determined by calculating the difference between the division angle (*θ*_D;_ measured at late anaphase) and *θ*_V_. The closer the absolute value of *θ*_D_-*θ*_V_ is to zero, the better the alignment between division orientation and cell shape (**Fig. 4A**).

**Fig. 4.**
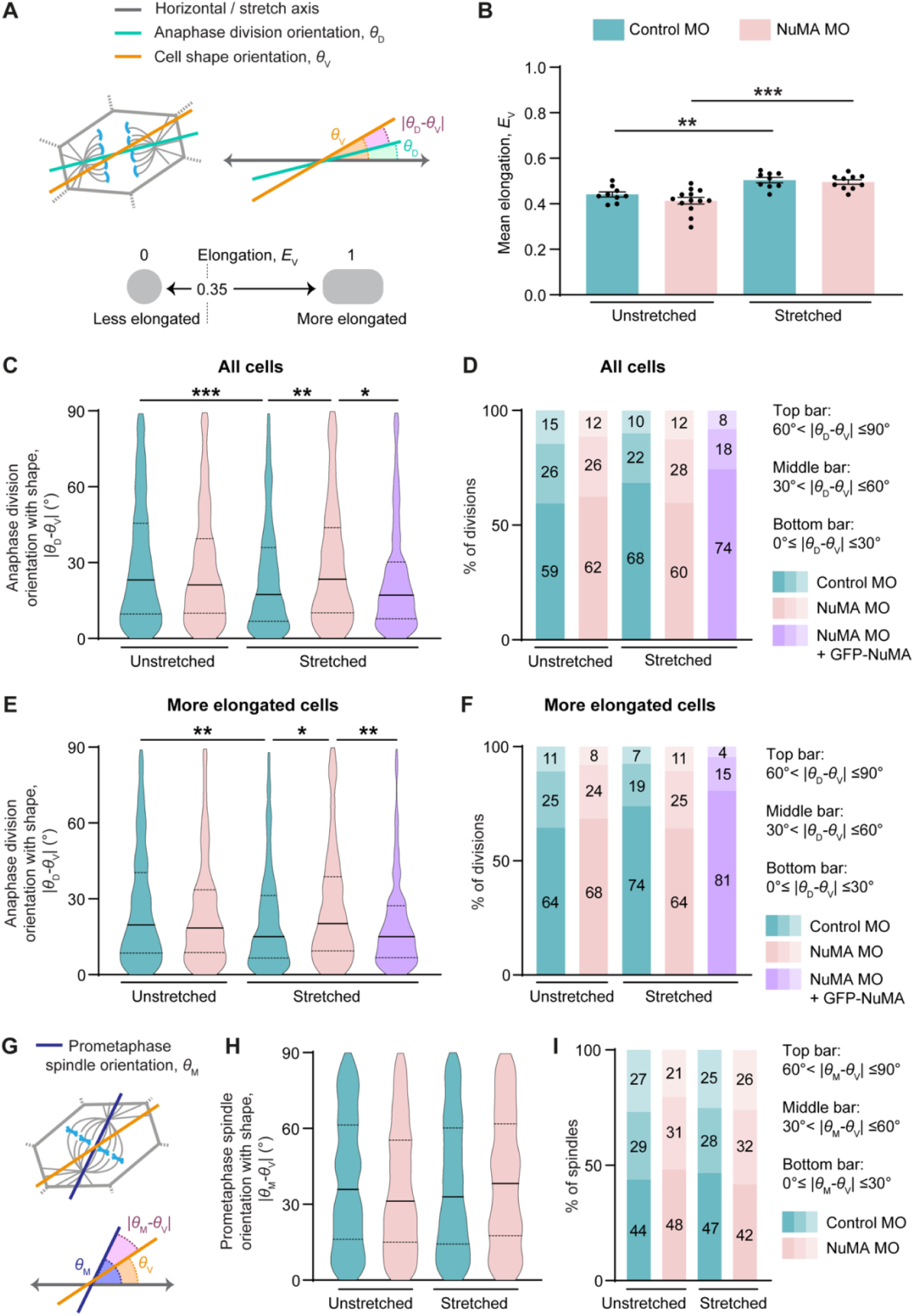
NuMA responds to anisotropic stretch to orient divisions with cell shape. **(A)** Schematic anaphase cell showing the eventual axis of division (*θ*_D_). The cell shape axis (*θ*_V_) is mathematically determined based on the position of the TCVs. The absolute value of the difference between *θ*_D_ and *θ*_V_ (|*θ*_D_*-θ*_V_|) is used as a measure of division alignment with cell shape. Elongation is mathematically determined based on the position of the TCVs (*E*_V_) and the value ranges from 0 to 1. Cells with *E*_V_ > 0.35 are classified as more elongated. **(B)** Mean elongation in unstretched and stretched control MO and NuMA MO tissues. Data were analysed using ordinary one-way ANOVA with Šidák’s multiple comparisons test. n=9 embryos; 13 embryos (unstretched, control MO and NuMA MO respectively); 9 embryos; 11 embryos (stretched, control MO and NuMA MO respectively). Error bars represent mean ± SEM. **(C)** Distribution of |*θ*_D_*-θ*_V_| in all cells of the unstretched and stretched control MO, NuMA MO, and rescue tissues. n=677 cells, 9 embryos; 609 cells, 13 embryos (unstretched, control MO and NuMA MO respectively); 608 cells, 9 embryos; 357 cells, 10 embryos (stretched, control MO and NuMA MO respectively); 304 cells, 11 embryos (rescue). **(D)** Percentage of divisions from each distribution in C binned into three partitions relative to cell shape. **(E)** Distribution of |*θ*_D_*-θ*_V_| in elongated cells of the unstretched and stretched control MO, NuMA MO, and rescue tissues. n=467 cells, 9 embryos; 411 cells, 13 embryos (unstretched, control MO and NuMA MO respectively); 489 cells, 9 embryos; 282 cells, 10 embryos (stretched, control MO and NuMA MO respectively); 238 cells, 11 embryos (rescue). **(F)** Percentage of divisions from each distribution in E binned into three partitions relative to cell shape. **(G)** Schematic prometaphase cell showing the spindle axis relative to the horizontal (*θ*_M_). The absolute value of the difference between *θ*_M_ and *θ*_V_ (|*θ*_M_*-θ*_V_|) is used as a measure of spindle alignment with cell shape at prometaphase. **(H)** Distribution of |*θ*_M_*-θ*_V_| in all cells of the unstretched and stretched control MO and NuMA MO tissues. n=647 cells, 9 embryos; 557 cells, 13 embryos (unstretched, control MO and NuMA MO respectively); 594 cells, 9 embryos; 331 cells, 10 embryos (stretched, control MO and NuMA MO respectively). Data in C, E, and H were analysed using Kruskal-Wallis test and Dunn’s multiple comparisons test. For C, E, and H, solid and dashed lines represent median and quartiles respectively. **(I)** Percentage of spindles from each distribution in H binned into three partitions relative to cell shape. Statistical significance: p<0.001***, p<0.01**, p<0.05*.

Hertwig’s rule dictates that most cells tend to orient their divisions with their long axis of shape (Hertwig, 1884). This was observed in both the unstretched and stretched tissues (**Fig. 4C**). In unstretched tissues, divisions were similarly aligned with cell shape in both control and NuMA knockdown tissues with no significant difference between the two distributions (**Fig. 4C**). Importantly, the application of stretch significantly improved division alignment with cell shape (|*θ*_D_-*θ*_V_| moves closer to 0°) in the control tissues (**Fig. 4C**). However, this improvement in division alignment with cell shape was not observed in the NuMA knockdown tissues despite a stretch (**Fig. 4C**). Therefore, there was a significant difference in the distribution of division orientation with cell shape between stretched control and stretched NuMA knockdown tissues.

To further probe the extent to which divisions were aligned with shape, we tripartite binned each distribution relative to the axis of cell shape (**Fig. 4D**). This showed that the majority of divisions aligned within 30° of the cell shape-axis in all conditions. In the unstretched control and NuMA knockdown tissues this was very similar with 59% and 62% divisions orienting closest to cell shape, respectively. Stretch enhanced this alignment to 68% divisions in the control tissues. However, this was lower at 60% in the stretched NuMA knockdown tissues, similar to that in unstretched NuMA knockdown tissues (**Fig. 4D**). Crucially, the reduction in alignment of division orientation with cell shape observed in the stretched NuMA knockdown tissues could be rescued by introducing GFP-NuMA over the endogenous knockdown background (**Fig. 4C, D**).

Importantly, the misalignment of divisions with cell shape in the stretched NuMA knockdown tissues was a robust phenotype since this was observed regardless of whether cell shape was measured based on TCVs, cell perimeter or cell area (**Fig. 4C; Fig. S4A, B**). We also investigated whether cell size, as measured by apical cell area, was a factor in the misorientation of divisions with cell shape in the stretched NuMA knockdown tissues. We found no significant effect of NuMA knockdown on cell area in either unstretched or stretched tissues (**Fig. S4C**). Additionally, there was no correlation between cell area and the alignment of division orientation with cell shape in any of the conditions (**Fig. S4D**).

Together, these data indicate that the alignment of the spindle with cell shape is significantly improved upon uniaxial stretch and that this process requires NuMA. Importantly, knockdown of NuMA does not reduce the alignment of division with cell shape in unstretched tissues. This suggests that NuMA is functioning to improve division alignment with cell shape but only in response to uniaxial stretch. However, a confounding factor here is that uniaxial stretch also changes the shape of cells, making them more anisotropic in shape (**Fig. 4A, B**). Since spindles are known to align more accurately with anisotropic cell shapes (Nestor-Bergmann et al., 2019) this increased anisotropy could also explain the difference in NuMA knockdown phenotype between stretched and unstretched tissues. We therefore sought next to exclude, as far as possible, the confounding factor of shape anisotropy.

### NuMA links anisotropic stretch to division orientation independently of cell shape

To exclude the confounding factor of shape anisotropy and address whether NuMA responds to force or cell shape, we computationally fractionated the whole population of cells into different shape subtypes based on cell elongation. In this way, we could compare cells of a similar shape across the different conditions. Elongation was mathematically determined based on the position of the TCVs (*E*_V_) and ranged from 0 to 1 (the closer the value is to 1, the more elongated the cell) (**Fig. 4A**). We focussed on the most elongated cells from each condition, since it is with these most anisotropically shaped cells that the most accurate measurement of the long axis is achieved. To classify a cell as either more or less elongated, we employed an *E*_V_ cutoff value of 0.35 such that cells with *E*_V_ > 0.35 were considered more elongated (Nestor-Bergmann et al., 2019) (**Fig. 4A**).

In these more elongated cells, division orientation with cell shape mirrored the effects seen in the overall cell population (**Fig. 4C, E**). In all cases, divisions were majorly oriented with cell shape with no significant difference between the unstretched control and NuMA knockdown tissues (**Fig. 4E, F**). Stretch improved division alignment with cell shape in the control tissues with 74% divisions orienting within 30° of the cell shape axis compared to 64% in the unstretched tissues (**Fig. 4F**). This improvement was negated in the stretched NuMA knockdown tissues but could be rescued by exogenous NuMA (**Fig. 4E, F**). Despite all the cells being of a similar shape anisotropy, the division misalignment with cell shape was only observed in the NuMA knockdown tissues specifically in the presence of a tissue stretch. This suggests that NuMA links spindle orientation to mechanical force directly rather than indirectly via cell shape.

Together, these data demonstrate that NuMA functions in mechanosensitive division orientation to finesse the ability of the spindle to find the long axis and therefore align with an axis of stretch. Moreover, this finessing by NuMA appears to be a direct response to mechanical force rather than an indirect response to a change in cell shape anisotropy.

### Mechanosensitive spindle reorientation with cell shape occurs later in mitosis

We next sought to determine whether NuMA-mediated mechanosensitive finessing of spindle orientation with cell shape occurs at a particular time during mitosis. Additionally, given that cells generally align divisions with the long axis of interphase shape, it is important to determine whether spindle alignment with interphase shape is determined early in mitosis. The finessing of division alignment with cell shape that we observe in stretched control tissues was measured at the end of metaphase (**Fig. 4C, D**). This coincides with our observation that NuMA is significantly enriched at the polar cortex at late metaphase in stretched tissues (**Fig. 1; Fig. S1**). To check whether this is coincidental, we asked whether a similar stretch-induced fine-tuning of spindle alignment with cell shape could be observed early in mitosis, at prometaphase, when there is no significant enrichment of NuMA at the polar cortex (**Fig. 1; Fig. S1**).

To do this, we measured the axis of the spindle relative to the horizontal/stretch axis at prometaphase (*θ*_M_) and used this to determine the absolute difference between spindle alignment and cell shape (|*θ*_M_-*θ*_V_|) at prometaphase (**Fig. 4G**). In contrast to the anaphase spindles that were predominantly oriented with cell shape (**Fig. 4C, D**), the spindles at prometaphase were less oriented with shape (**Fig. 4H, I**). This was the case for both control and NuMA knockdown tissues. For example, in the unstretched control tissues, 59% of spindles at anaphase were oriented within 30° of cell shape (**Fig. 4D**) whereas fewer spindles (44%) were oriented to this extent at prometaphase (**Fig. 4I**). This suggests that even though spindle alignment with cell shape is not entirely random at the onset of mitosis, there is further enhancement of division alignment with shape later in mitosis.

Interestingly, in contrast to the anaphase spindles (**Fig. 4C, D**), application of a tissue stretch did not improve spindle alignment with cell shape at prometaphase in the control tissues (**Fig. 4H, I**). The percentage of spindles aligning within 30° of the cell shape-axis remained similar at 44% and 47% for unstretched and stretched control tissues respectively (**Fig. 4I**). In addition, there was no misalignment upon NuMA knockdown (**Fig. 4H, I**).

Together, these data indicate that spindle orientation along the long axis of interphase shape is not strictly determined at the onset of mitosis but is rather modulated throughout mitosis. Furthermore, external mechanical forces continue to finesse this spindle orientation during metaphase via a dynamic increase in NuMA recruitment to the polar cortex in stretched tissue.

## Discussion

The fundamental mechanisms of cell division orientation are well characterised but considerably less is known about how mechanical forces are sensed by the mitotic spindle to orient divisions. In this work, we identify NuMA as a key link between external forces and spindle orientation. We show that NuMA has a mechanosensitive cortical localisation, with earlier mitotic recruitment to the polar cortex in tissues under uniaxial stretch. This early recruitment is shown to have a direct impact on spindle dynamics, increasing mitotic spindle oscillation and, therefore, increasing cortical force generation to position the spindle. Knockdown of NuMA reduces spindle oscillation in stretched tissues and reduces the accuracy of spindle alignment with a global axis of tension and with local cell shape. We therefore suggest that NuMA provides a critical link between anisotropic tension and the spindle to precisely orient cell divisions.

A significant proportion of our knowledge of spindle orientation mechanisms stems from fundamental studies in *Drosophila* (Bosveld et al., 2016; Bowman et al., 2006; Izumi et al., 2006; Siller et al., 2006). For example, the *Drosophila* homologue of NuMA, Mud, localises at TCJs to mediate spindle alignment with interphase cell shape in the pupal notum tissue (Bosveld et al., 2016). We show that, unlike Mud, NuMA is not enriched at the TCJs in the *Xenopus* animal cap but is rather distributed around the cell edges (**Fig. 1; Fig. S1**). Our previous work has suggested that LGN localisation may be more akin to Mud in vertebrate tissue, being enriched at cell vertices and important for shape-dependent spindle orientation (Nestor-Bergmann et al., 2019). The differing localisation patterns of LGN and NuMA suggest, as has been shown in other systems, that NuMA may be targeted to the cortex independently of LGN (Bosveld et al., 2016). We speculate that the differential localisation of NuMA and LGN may be indicative of a layering of distinct mechanisms to efficiently orient the spindle with anisotropic tension: shape-mediated alignment by LGN and force-mediated alignment by NuMA. This layering of mechanisms provides robust and accurate division orientation.

We propose that the earlier recruitment of NuMA to the polar cortex seen when we stretch the tissue provides a mechanism to directly finesse spindle alignment with external force. Indeed, when NuMA levels are depleted, we observe a misalignment of divisions with the global axis of stretch, similar to the misalignment seen in a previous study on fixed keratinocytes (Seldin et al., 2013). Crucially, this earlier recruitment of NuMA in stretched tissues coincides with spindles becoming more dynamic and undergoing an increase in oscillatory movement. Mathematical modelling has shown that increases in oscillatory movement reflect an increase in the number of cortical force generators (*N)* pulling on the astral microtubules to position the spindle (Grill et al., 2005; Hargreaves et al., 2024). When a tissue is stretched, we see an earlier recruitment of NuMA and a concomitant increase in spindle oscillation, indicating increased cortical force generation to facilitate more accurate spindle positioning. Significantly, what we see in stretched tissue is an acceleration of what usually occurs in unstretched tissue: NuMA levels at the polar cortex increase as mitosis progresses to reach a maximum in anaphase, with a corresponding increase in spindle oscillation and, therefore, cortical force generation. This progressive focusing of NuMA localisation helps hone spindle position and also allows continual adjustment to the external environment. Indeed, building on previous work in single cells (Fink et al., 2011), we show that spindle alignment with cell shape is not strictly determined at the onset of mitosis, but rather finessed by external force during metaphase, with a direct role for NuMA in this process.

Significantly, our results indicate that NuMA links spindle orientation directly to force, rather than indirectly via force-induced changes in cell shape. Since cell shape will reflect local forces within the heterogeneous epithelial layer (Nestor-Bergmann et al., 2018; Nestor-Bergmann et al., 2019), separating mechanisms reliant on cell shape from those involving direct force-sensing is challenging. To overcome this issue, we filtered cells according to cell shape and compared cells that were the same shape in unstretched and stretched tissue. We chose to concentrate on the most elongated cells as the shape of these cells can be most accurately measured. Crucially, we saw spindle alignment with cell shape improve when tissue is stretched, even in equivalently shaped cells, and we saw a loss of this stretch-induced improvement when NuMA was knocked down. Seeing this tension-dependent effect in elongated cells is interesting as, in other tissue models, the extent of cell elongation often overrides any direct force-sensing mechanism (Campinho et al., 2013; Godard et al., 2020; Mao et al., 2013; Scarpa et al., 2018; Tang et al., 2018; Wyatt et al., 2015). Our data shares interesting parallels with findings from the *Drosophila* pupal notum where cells in the midline, despite being the most elongated, are less efficient at orienting divisions with cell shape under low tissue tension (Lam et al., 2020). In these elongated cells, fewer spindles undergo Mud-dependent rotations toward the long axis compared to cells outside the midline under higher tissue tension. Additionally, upon increasing tension within the midline, the elongated cells become more efficient at orienting divisions with shape (Lam et al., 2020).

An important remaining question is what facilitates NuMA’s earlier recruitment to the polar cortex in stretched tissues? Our findings of a mechanosensitive localisation of NuMA are reinforced by studies where cells on adhesive micropatterns localise NuMA to cortical regions that are in direct contact with the micropattern (Machicoane et al., 2014). Moreover, cortical distance-sensing mechanisms such as exclusion of NuMA from the equatorial cortex by a Ran-GTP gradient (Kiyomitsu and Cheeseman, 2012) and from the polar cortex by polo-like kinase 1 (PLK1) (Sana et al., 2018) could be at play here. Interestingly, NuMA can be localised to the cortex dependent on its state of phosphorylation (Gallini et al., 2016; Kotak et al., 2013; Matsumura et al., 2012; Zheng et al., 2014) and interactions with actin-binding proteins may further facilitate NuMA/LGN localisation to the cortex (Carminati et al., 2016; Kiyomitsu and Cheeseman, 2013; Machicoane et al., 2014; Sharma et al., 2020). Intriguingly, NuMA also contains a large coiled-coil region (Compton et al., 1992) and, in other proteins, coiled-coils have been shown to be mechanosensitive (Craven et al., 2022; Premchandar et al., 2016). Therefore, it is interesting to speculate whether NuMA’s phosphorylation state, interaction with actin-binding proteins and/or coiled-coil domain have a role in localising NuMA earlier to the polar cortex in stretched tissues and these will provide important avenues for future study.

In conclusion, our findings support a model whereby anisotropic tissue tension drives earlier recruitment of NuMA to the polar cortex during mitosis. This enhanced cortical localisation of NuMA modulates increased spindle oscillations and, therefore, increased cortical force generation to pull the spindle into position. Ultimately, NuMA mediates a dynamic, force-dependent, finessing of spindle alignment with anisotropic tension to ensure accurate cell division orientation.

## Materials and methods

### DNA constructs and morpholino oligonucleotides

pCS2+/mCherry-Histone2B (Kanda et al., 1998), pCS2+/GFP-α-tubulin (Woolner et al., 2008), pCS2+/BFP-CAAX (Larson and Bement, 2017), and pCS2+/GFP (Burkel et al., 2007) were gifts from Bill Bement (University of Wisconsin-Madison, US). pCS2+/GFP-CAAX (Reyes et al., 2014) was a gift from Ann Miller (University of Michigan, US). pCS2+/GFP-NuMA was generated by excising the NuMA sequence from pEGFP/C1-NuMA (Addgene, 81029) and ligating into Bgl2 and EcoR1 sites in pCS2+/GFP. Morpholino (MO) sequences (Gene Tools, LLC) used were as follows: Standard Control (5’-CCTCTTACCTCAGTTACAATTTATA-3’), NuMA1.S (5’-GTCATTATGCTTCAGCACTTCTCCC-3’), and NuMA1.L (5’-GCCATTTTGTTTCACTACTTTTCCC-3’).

### *Xenopus laevis* ovulation induction and natural mating

*Xenopus laevis* were housed within tanks maintained by the in-house animal facility at the University of Manchester. The female and male frogs were used for embryo collection and natural mating respectively. Females were pre-primed 4-7 days in advance of egg collection with 50 U of pregnant mare’s serum gonadotrophin (MSD Animal Health) injected into the dorsal lymph sac. Before the day of egg collection, females and males were primed with 200 U and 100 U of human chorionic gonadotrophin, Chorulon (MSD Animal Health) respectively, injected into the dorsal lymph sac. The primed frogs were maintained in individual tanks containing aquarium water, and 16 to 18 hours later, a female and male frog were transferred to the same tank for natural mating. Eggs were collected from the tank 2-5 hours later. All *Xenopus* work was performed using protocols approved by the UK Government Home Office and covered by Home Office Project Licences PFDA14F2D (Licence holder: Enrique Amaya) and PP1859264 (Licence holder: Karel Dorey) as well as Home Office Personal Licences held by N. Tarannum, D. Hargreaves, G. Goddard, and S. Woolner.

### Embryo processing and microinjection

Embryos collected from natural mating were processed as previously described (Goddard et al., 2020; Woolner and Papalopulu, 2012). mRNA was synthesised as described previously (Sokac et al., 2003) and each cell was injected with a total volume of 4.2 nl or 2.1 nl for injections at 2-cell or 4-cell stages respectively. RNA was microinjected into both cells at 2-cell stage at the following needle concentrations: 0.1 mg/ml mCherry-Histone2B, 0.1 mg/ml BFP-CAAX, 0.1 mg/ml GFP-CAAX, 0.5 mg/ml GFP-α-tubulin, 0.25 mg/ml GFP-human NuMA (overexpression experiments), 0.01 mg/ml GFP-NuMA (morphant rescue experiments). For NuMA knockdown/rescue and metaphase plate tracking experiments, MOs prepared as 1 mM stocks were microinjected at a needle concentration of 1 mM. NuMA1.S MO and NuMA 1.L MO were injected into all cells at 2-cell and 4-cell stages respectively. Control MO was injected similarly for respective control experiments. Following microinjection, embryos were incubated in 0.1% MMR at 16°C for approximately 20 hours.

### Animal cap dissection and culture

Animal cap tissue was dissected from embryos at stage 10 of development (early gastrula stage) as previously described (Joshi and Davidson, 2010). Dissected animal caps were cultured *ex vivo* in Danilchik’s for Amy explant culture media (53 mM NaCl, 5 mM Na_2_CO_3_, 4.5 mM Potassium gluconate, 32 mM Sodium gluconate, 1 mM CaCl_2_, 1 mM MgSO_4_, pH 8.3 with Bicine) with 0.1% BSA (Sigma, A7906) on a fibronectin-coated PDMS membrane chamber which was prepared as described previously (Goddard et al., 2020). Explants were held in place by a coverslip fragment and allowed to adhere to the fibronectin at 18°C for 2 hours before imaging (Goddard et al., 2020).

### Animal cap stretch manipulation and imaging

The PDMS membrane chamber was placed on a stretch apparatus (custom-made by Deben UK Ltd.) fixed securely to the stage of the confocal microscope and then subjected to one of the following stretch manipulations: unstretched (0.5 mm displacement to remove sag on the membrane) or uniaxial stretch (8.6 mm displacement of one arm). Images were acquired with the Leica Application Suite X (LAS X) software using hybrid detectors (HyDs) in a 20-23°C environment.

For GFP-NuMA localisation, NuMA knockdown (except for metaphase plate tracking), and rescue experiments, images were collected on a Leica TCS SP8 AOBS upright confocal. GFP-NuMA localisation and NuMA knockdown/rescue images were acquired using a 63x/0.9 HCX Apo U-V (dipping lens) and a 20x/0.50 HCX Apo U-V-I (dipping lens) respectively with the following settings: 1 AU pinhole, 400 Hz bidirectional scanning, 1X confocal zoom, 512 x 512 pixel resolution. Z stacks were collected with an optical section of 2 μm between steps and a time interval of 30 s between frames for up to 2.5 hours. For metaphase plate tracking experiments, images were collected on a Leica Stellaris 8 upright confocal using a 20x/0.50 HCX Apo U-V-I (dipping lens) and the following settings: 1.9 AU, 600 Hz bidirectional scanning, 1.5X confocal zoom, 1024 x 1024 pixel resolution. Z stacks were collected with an optical section of 10 μm between steps and a time interval of 5 s between frames for up to 2.5 hours. GFP-NuMA localisation images were collected at a bit depth of 16 bits. All other images were collected at 8 bits. Maximum intensity projections of these 3D stacks are shown in the results.

### Immunofluorescence

Immunofluorescence was performed as previously described (Moruzzi et al., 2021). Stage 10 embryos injected with control MO and NuMA MO were fixed and incubated in primary and secondary antibodies in 1X Tris-buffered saline with Nonidet P-40 (TBSN) (155 mM NaCl, 10 mM Tris-HCl pH 7.4, 0.1% Nonidet P-40) containing 10 mg/ml BSA at the following dilutions: mouse anti-α-tubulin (Sigma, T9026) (primary; 1:200) and goat anti-mouse Alexa Fluor 488 (Thermofisher, A-11001) (secondary; 1:400). Embryos were also incubated with 10 µg/ml DAPI (Invitrogen, D2149). Samples were methanol dehydrated and then cleared and mounted in Murray’s Clear (2:1 benzyl benzoate:benzyl alcohol). Images were collected on a Leica TCS SP8 AOBS inverted confocal containing the LAS X software using HyDs. Images were acquired using a 40x/1.30 HC PL APO CS2 (oil) objective with the following settings: 1 AU pinhole, 400 Hz bidirectional scanning, 0.75X confocal zoom, 1024 x 1024 pixel resolution. Z stacks were collected with an optical section of 1.5 μm between steps. Maximum intensity projections of these 3D stacks are shown in the results.

### Whole embryo imaging

To show that the metaphase plate can be used as a proxy for spindle movements, embryos at stage 10-11 were mounted as described previously (Woolner et al., 2010). Single focal plane live-cell images of spindles were collected at room temperature every 20 s using a FluoView FV1000 Olympus microscope with FluoView acquisition software (Olympus) and a 60x, 1.35 NA U Plan S Apochromat objective.

### Morphant embryo survival

Uninjected, control MO, and NuMA MO-injected embryos in 0.1% MMR were imaged during early development. Embryos were imaged on a Zeiss Lumar V.12 Stereoscope using the Zen Pro software. Images were acquired every 10 minutes at 6.4X zoom for up to 48 hours in a 20-22°C environment. The percentage of dead embryos were determined for each independent experiment.

### Western blotting

Western blotting was performed as described previously (Moruzzi et al., 2021). Stage 10 embryos injected with NuMA MO, control MO, and NuMA MO with GFP-NuMA were lysed and fractionated by SDS-PAGE on 4-15% Mini-PROTEAN TGX Stain-Free Protein Gels (Bio-rad, 4568083) at 120 V. Proteins were transferred to a 0.45 μm nitrocellulose membrane (GE Healthcare, 10600002) at 30 V for 16 hours at 4°C. The membrane was blocked by incubation with 5% non-fat milk in 1X Tris-buffered saline with tween (TBST) (10 mM Tris pH 8.0, 150 mM NaCl, 0.5% tween 20) and incubated with the following antibodies in blocking solution: rabbit anti-NuMA1 (1:500) (Prestige Antibodies, HPA029912), mouse anti-α-tubulin (1:1000), goat anti-rabbit IgG H&L IRDye 800CW (1:5000) (Abcam, ab216773), and donkey anti-mouse IgG H&L IRDye 680RD (1:5000) (Abcam, ab216778). Membranes were imaged on the Odyssey CLX LI-COR. Band intensity was measured using Image Studio (LI-COR Biosciences). The intensity of each band was normalised to background fluorescence. The band for NuMA was then normalised to α-tubulin. The fold change relative to the control MO was then calculated.

### Image processing and quantification

Image processing and analysis were performed using ImageJ (Fiji) (Schindelin et al., 2012; Schneider et al., 2012). Cortical localisation analysis was performed on sum intensity projections. All other analyses were performed on maximum intensity projections.

#### Cortical localisation

The cortical fluorescence of GFP-NuMA was measured by drawing ROIs at the following regions of the cortex: TCVs (oval ROI), cell edges (polygon ROI), entire cortex (polygon ROI), polar and equatorial cortex (oval and polygon ROI). To minimise bias, all ROIs were drawn while visualising the BFP-CAAX channel. Measurements were taken at different stages of mitosis including prometaphase (onset of visualising condensed chromosomes), mid metaphase (halfway between prometaphase and late metaphase), late metaphase (30 seconds before the onset of anaphase) and early anaphase (30 seconds after the onset of anaphase). Fluorescence measurements were avoided at mid anaphase due to the close proximity of spindle pole NuMA and cortical NuMA. Five background fluorescence measurements were taken per embryo, and the grand mean of the background fluorescence was used to calculate the corrected total fluorescence (CTF) density of GFP-NuMA using the equation

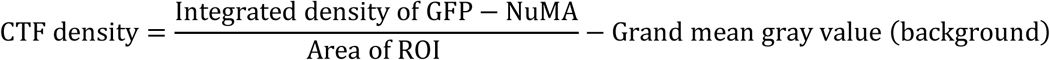

For every mitotic stage, two values of CTF density were obtained for the polar cortex and the equatorial cortex. The values were then averaged for each cortical region. CTF density values for TCVs and cell edges were also averaged based on the number of TCVs and edges respectively. The mean CTF density values are shown in the results. The fold change of polar cortical fluorescence was calculated relative to the equatorial cortical fluorescence and the fold change of TCVs was calculated relative to the cell edges. The fold change values are shown in the results.

#### Metaphase plate/spindle tracking

The metaphase plate was used as a proxy to track spindle movements during metaphase. Each end of the metaphase plate was tracked through time using the “MTrackJ” plugin (Meijering et al., 2012). For every time point, *x* and *y* coordinates were obtained for both ends of the metaphase plate. The coordinates were processed through an in-house Julia code to generate a line connecting both ends of the metaphase plate. The angle between this line and the horizontal at each time point was calculated (*θ*_plate_) and used to determine the angle of the mitotic spindle (*θ*_M_=*θ*_plate_ - 90°).

The overall spindle trend (*θ*_T_) was determined using a moving median in a sliding window of width Δ*t*=125 s. The median at the beginning and end of the signal time bounds were determined by reflecting the signal through the boundary. The root mean squared (RMS) of the angle change was determined by

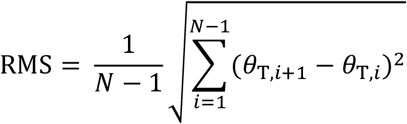

where *i* indicates the timepoint of *θ*_T,*i*_, where the signal is measured for *N* sequential timepoints.

#### Measuring oscillations

An autocorrelation function (ACF) was used to determine whether de-trended spindle angles were oscillatory, as well as their associated period of oscillation. Peaks above threshold in the autocorrelation of the signal at increasing time-lags were used to identify an oscillatory spindle, with the associated time-lag corresponding to the oscillation period. The threshold was defined as

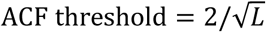

for *L* the length of the input signal (the number of frames of measured spindle angles, from prometaphase to anaphase).

Of those deemed oscillatory, the range of oscillation was determined as the range between the upper and lower bounds of the signal, omitting outliers. Outliers were omitted using the interquartile range (IQR) of the signal, and the lower quartile (Q1) and upper quartile ( Q3) thresholds, such that outliers were points below a lower bound

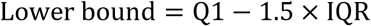

or above the upper bound

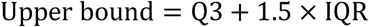

#### Division orientation

Division orientation was measured using the “straight-line” tool as described previously (Nestor-Bergmann et al., 2019). A line was drawn between the two separating daughter nuclei of a cell dividing in-plane at late anaphase. Using the ROI manager, the angle of division relative to stretch (horizontal axis) (*θ*_D_) was calculated. The data were also separated into three 30° bins between 0 to 90° and the percentage of divisions within each bin was calculated.

#### Division rate

The first frame of each time-lapse experiment was extracted, and the number of nuclei was counted using the “cell counter” plugin. The division rate (percentage of total cells entering mitosis per hour) was calculated.

#### Spindle morphology

The number of bipolar and multipolar spindles was counted from immunofluorescence images using the “cell counter” plugin. The percentage of multipolar spindles was calculated for each embryo.

#### Single cell analysis of spindle alignment with cell shape

Single cells were traced as previously described (Nestor-Bergmann et al., 2019). Cell edges and junctions were manually traced 60 seconds before nuclear envelope breakdown using the “Paintbrush tool”. The frame number and *x* and *y* coordinates of the cell’s position were noted. The traces were processed using in-house python scripts (Nestor-Bergmann et al., 2019) to extract cell area, elongation (*E*), and the principal axis of cell shape (*θ*). *E* and *θ* were determined based on the position of TCVs (*E*_V_ and *θ*_V_), perimeter (*E*_P_ and *θ*_P_), and area (*E*_A_ and *θ*_A_). The orientation of cell division relative to cell shape was determined by calculating the difference between division orientation (*θ*_D_) and either *θ*_V_, *θ*_P_ or *θ*_A_. Cells were fractionated into either less or more elongated phenotypes based on *E*. *E* > 0.35 were classified as more elongated cells. This cut-off value was based on our previous work where we used circularity (1-*E*) to classify cell shape (Nestor-Bergmann et al., 2019).

For analysis of spindle alignment with cell shape at prometaphase, the prometaphase plate was used as a proxy for measurement of the spindle axis. Using the “straight-line” tool, a line was drawn between the ends of the prometaphase plate. The angle between this line and the horizontal was calculated (*θ*_plate_) and used to determine the angle of the spindle (*θ*_M_=*θ*_plate_ - 90°). The orientation of the prometaphase spindle relative to cell shape was determined by calculating the difference between the *θ*_M_ and *θ*_V_. The data for spindle orientation with cell shape were also separated into three 30° bins between 0 to 90° and the percentage of spindles within each bin was calculated.

### Data presentation and statistical analysis

Rose histograms were generated using in-house Python scripts (Nestor-Bergmann et al., 2019). All other graphs were plotted using GraphPad Prism 10 (GraphPad Software, San Diego, USA). Figures were prepared using Adobe Illustrator and Adobe Photoshop (Adobe, San Jose, USA). Statistical analysis was performed using GraphPad Prism 10. Samples were tested for normality using the D’Agostino-Pearson normality test and the Shapiro-Wilk normality test. Normality tests were passed at p>0.05. Following the assessment of normality, statistical tests were chosen as appropriate. For all statistical analyses, p values less than 0.05 were considered statistically significant. All statistical analysis and p values are reported in the figure legends.

### Online supplemental material

Four supplemental figures and seven videos are included. Fig. S1 shows the quantification of cortical NuMA at different regions of the cell cortex in stretched and unstretched tissues. Fig. S2 shows the modelling and measurement of spindle movements. Fig. S3 shows the effect of NuMA knockdown on embryonic development, spindle structure and cell division rate. Fig. S4 shows the effect of NuMA knockdown on apical cell area, division alignment with cell shape and the correlation between these two factors. Video 1 shows the localisation of NuMA in the animal cap explant. Video 2 shows the localisation of NuMA in a cell at interphase and mitosis. Video 3 shows a mitotic cell undergoing spindle oscillations. Video 4 shows a mitotic cell where the spindle does not oscillate. Video 5 shows cell divisions in a stretched animal cap explant. Video 6 shows a cell dividing along the axis of tissue stretch. Video 7 shows a cell dividing perpendicular to the axis of stretch.

## Acknowledgements

This work was supported by Wellcome Trust 4-year PhD Studentships for N.T. and D.H. (215210/Z/19/Z and 220054/Z/19/Z respectively), a Leverhulme Trust Research Project Grant (RPG-2021-394), a Wellcome Trust/Royal Society Sir Henry Dale Fellowship to S.W. (098390/Z/12/Z), and a Wellcome Trust Career Development Award to S.W. (225408/Z/22/Z). The Bioimaging Facility microscopes used in this study were purchased with grants from BBSRC, Wellcome, and the University of Manchester Strategic Fund. Thanks to Peter March, Roger Meadows, and Steve Marsden from the Biomaging team for their help with microscopy. Also, thanks to members of the Biological Services Facility for their help with the maintenance of *Xenopus laevis* colonies.

## Author contributions

S. Woolner, N. Tarannum, D. Hargreaves, and O.E. Jensen designed the study; N. Tarannum and D. Hargreaves developed methodology; N. Tarannum, D. Hargreaves, D. Ilieva, G. Goddard, and S. Woolner analysed the data; O.E. Jensen and S. Woolner acquired funding; N. Tarannum, D. Hargreaves, O.E. Jensen, and S. Woolner wrote the paper.

