## Supplemental information for "Mitotic spindle orientation and dynamics are fine-tuned by anisotropic tension via NuMA localisation"

### Figures

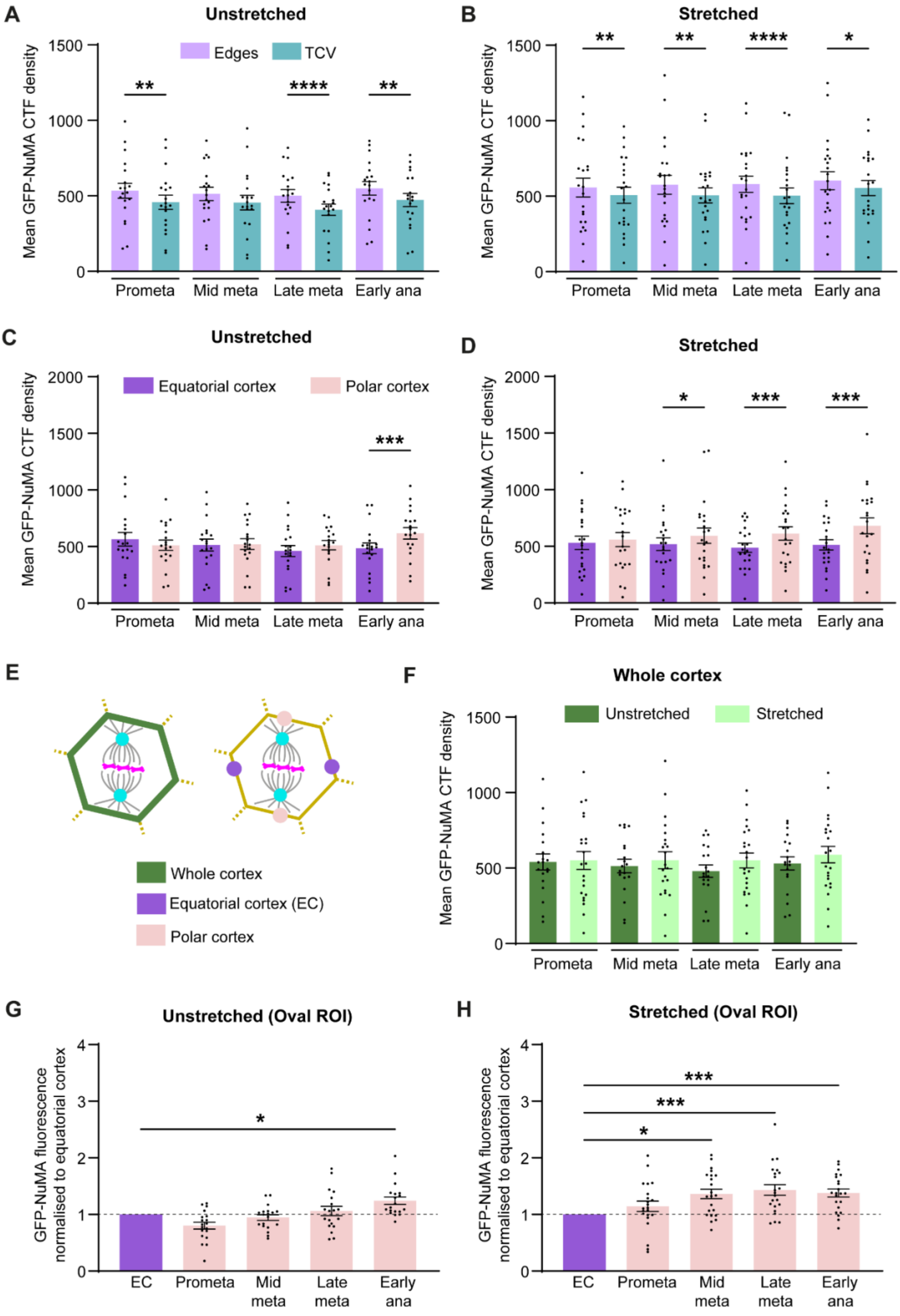
A
Unstretched
Edges
TCV
Mean GFP-NuMA CTF density
Prometa
Mid meta
Late meta
Early ana
B
Stretched
C
Unstretched
Equatorial cortex
Polar cortex
D
Stretched
E
Whole cortex
Equatorial cortex (EC)
Polar cortex
F
Whole cortex
Unstretched
Stretched
G
Unstretched (Oval ROI)
GFP-NuMA fluorescence normalised to equatorial cortex
EC
H
Stretched (Oval ROI)

**Fig. S1. Cortical localisation of NuMA at specific regions and over the whole cortex.**

**(A, B)** Mean corrected total fluorescence (CTF) density of GFP-NuMA at tricellular vertices and cell edges in unstretched and stretched tissues respectively. Respective normalised data are shown in Fig. 1C, D.

**(C, D)** Mean corrected total fluorescence (CTF) density of GFP-NuMA at polar cortex and equatorial cortex (polygon ROI) in unstretched and stretched tissues respectively. Respective normalised data are shown in Fig. 1E, F. Data in A, B, C, and D were analysed using RM one-way ANOVA with Geisser-Greenhouse correction and Šidák's multiple comparisons test.

**(E)** Fluorescence measurements of GFP-NuMA were taken throughout the entire cortex as well as polar and equatorial cortex during mitosis. Polar and equatorial cortex were measured using an oval ROI.

**(F)** Mean corrected total fluorescence (CTF) density of GFP-NuMA throughout the entire cortex in unstretched and stretched tissues. Data were analysed using ordinary one-way ANOVA with Šidák's multiple comparisons test.

**(G, H)** Mean normalised GFP-NuMA fluorescence at polar cortex relative to equatorial cortex (oval ROI) in unstretched and stretched tissues respectively. Data were analysed using Friedman test and Dunn's multiple comparisons test. For all data, n=19 cells, 5 embryos (unstretched); 22 cells, 6 embryos (stretched). Statistical significance: $p<0.0001$****, $p<0.001$***, $p<0.01$**, $p<0.05$*. All error bars represent mean ± SEM.

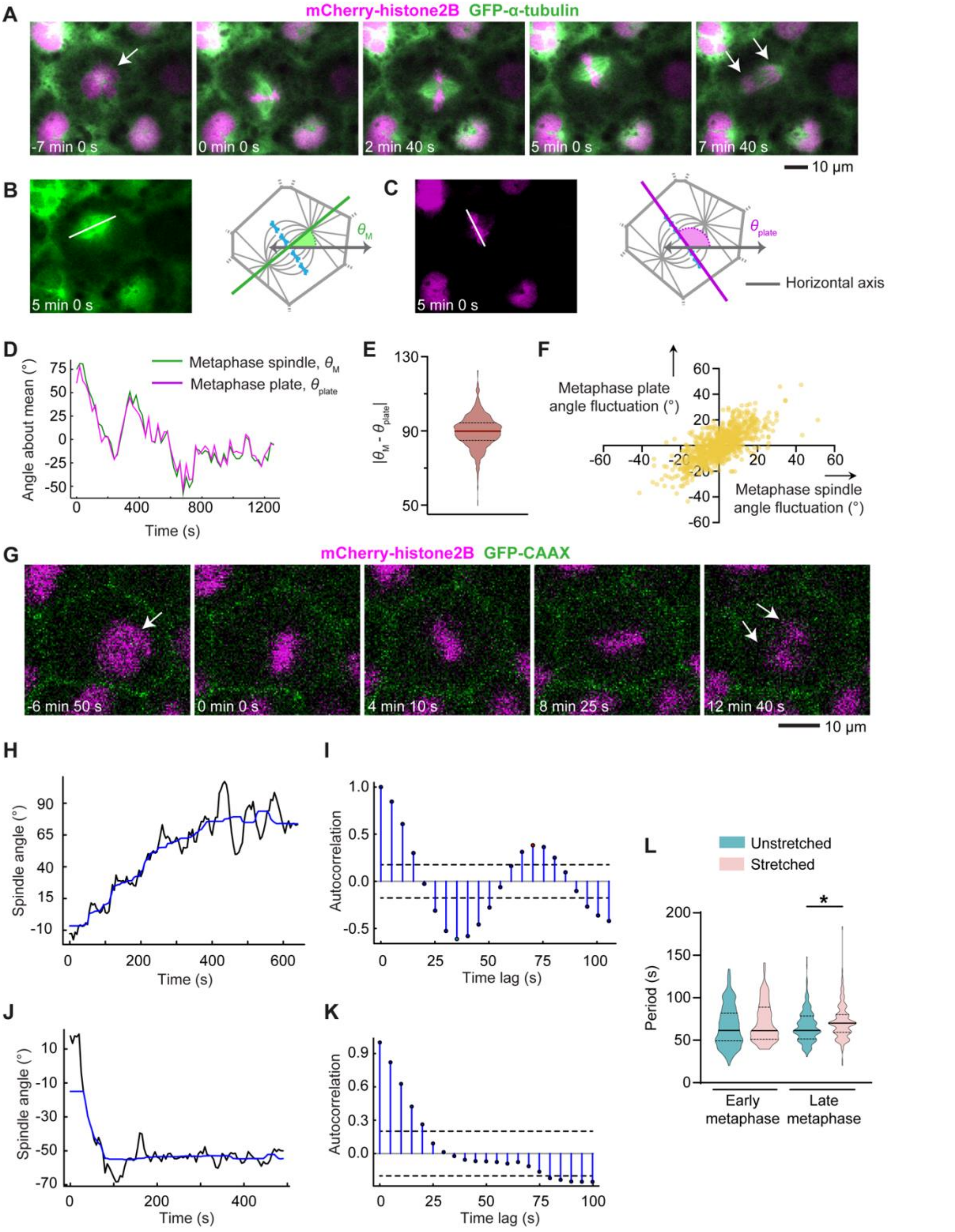


**Fig. S2. Metaphase plate movements can be used as a proxy for spindle movements.**

**(A)** A representative cell at interphase and mitosis in the embryo expressing a microtubule marker (GFP-α-tubulin; green) and chromatin marker (mCherry-histone2B; magenta). White arrow in the first frame identifies the dividing cell. White arrows in the final frame indicate the separation of chromosomes at anaphase. Time is chosen such that 0 minute coincides with the formation of the metaphase plate.

**(B, C)** Schematics of angle measurements for (B) the mitotic spindle ($\theta_M$), and (C) the metaphase plate ($\theta_{plate}$).

**(D)** Example measurements of the spindle and metaphase plate angles in time, normalised about their respective means.

**(E)** Angle difference between metaphase plate and the spindle. Data were analysed using the Wilcoxon signed rank test compared with a median of 90º.

**(F)** Inter-frame angle displacements for metaphase plate and spindle angle measurements. Data analysed using the Spearman rank correlation test: $p<0.0001$. For E and F, n=901 individual measurements from 15 cells.

**(G)** A representative cell at interphase and mitosis in a stretched tissue expressing a membrane marker (GFP-CAAX; green), chromatin marker (mCherry-histone2B; magenta), and containing control morpholino. White arrow in the first frame identifies the dividing cell. White arrows in the final frame indicate the separation of chromosomes at anaphase. Time is chosen such that 0 minute coincides with the formation of the metaphase plate. Related to Video 3.

**(H, J)** Example spindle angles undergoing rotational and (H) oscillatory and (I) non-oscillatory movements from prometaphase until the start of anaphase. The overall spindle movement (blue line) is determined using a moving median in a temporal window of $\Delta t$=125 s. H and J are related to Video 3 and 4 respectively.

**(I, K)** Autocorrelation functions for spindle angles timeseries H and J respectively. Dashed lines indicate significance thresholds. Significant peaks denoting an oscillatory signal in I; K shows a non-oscillatory signal.

**(L)** Period of oscillation for oscillatory spindles in stretched and unstretched tissues, in both early and late metaphase. Data were analysed using a mixed-effects model and Fisher's least significant difference test: $p<0.05$*. n=152 cells, 8 embryos; 193 cells, 8 embryos (unstretched, early and late metaphase respectively); 156 cells, 8 embryos; 174 cells, 8 embryos (stretched, early and late metaphase respectively). For H and L, solid and dashed lines represent median and quartiles respectively.

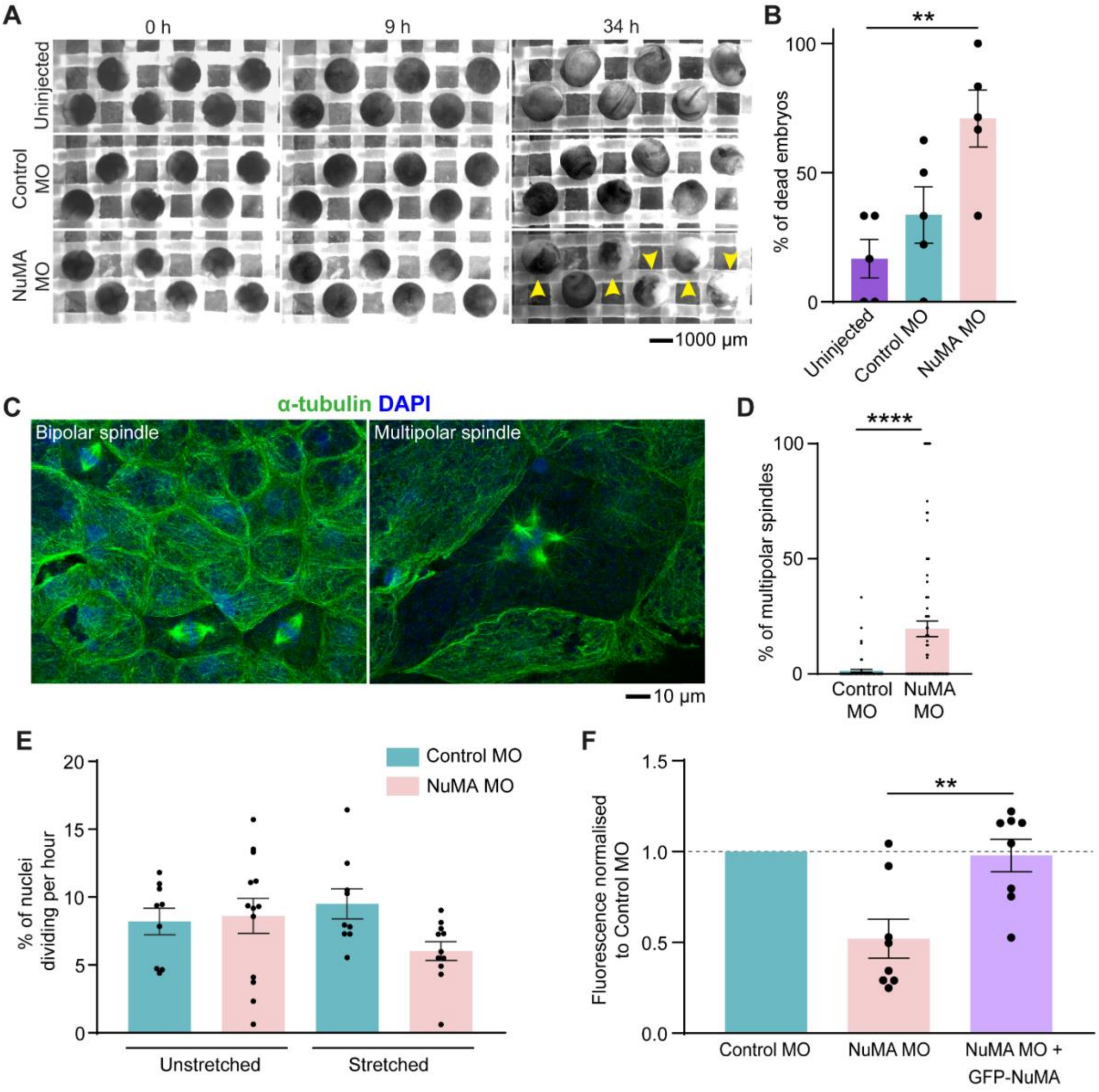


**Fig. S3. Characterisation of the effect of NuMA knockdown on embryonic development, spindle structure and cell division rate.**

**(A)** Uninjected embryos and embryos injected with control MO and NuMA MO during early development. 9 h timepoint corresponds to stage 10 (early gastrula) embryos. Yellow arrows indicate dead embryos.

**(B)** Percentage of dead embryos for uninjected, control MO-injected, and NuMA MO-injected embryos. Data were analysed using one-way ANOVA with Tukey's multiple comparison's test. n=5 independent repeats per condition from 37 embryos (uninjected); 35 embryos (control MO); 30 embryos (NuMA MO).

**(C)** Fixed embryos stained to visualise microtubules (α-tubulin, green) and DNA (DAPI, blue) showing bipolar and multipolar spindles.

**(D)** Percentage of multipolar spindles in control MO-injected and NuMA MO-injected fixed embryos. Data were analysed using Kolmogorov-Smirnov test. n=35 embryos from 743 cells (control MO); 42 embryos from 486 cells (NuMA MO).

**(E)** Division rate in control MO and NuMA MO tissues. Data were analysed using ordinary one-way ANOVA and Šidák's multiple comparisons test. n=9 embryos (unstretched control MO); 13 embryos (unstretched NuMA MO); 9 embryos (stretched control MO); 11 embryos (stretched NuMA MO).

**(F)** NuMA expression following knockdown of NuMA and addition of exogenous GFP-NuMA normalised to control MO. Data were analysed using Kruskal-Wallis test and Dunn's multiple comparisons test. n=8 independent repeats per condition from 80 embryos. Statistical significance: $p<0.0001$****, $p<0.01$**. All error bars represent mean ± SEM.

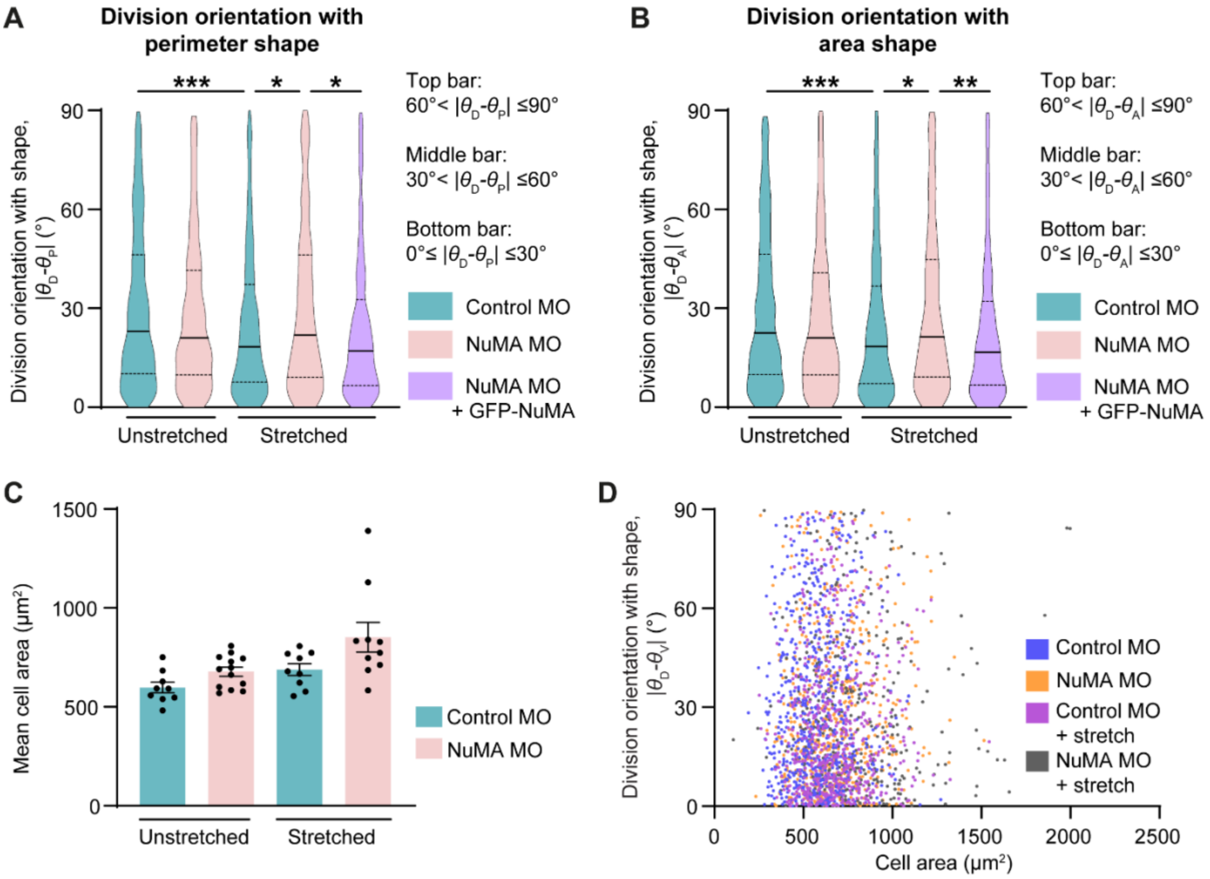


**Fig. S4. Effect of NuMA knockdown on apical cell area and division orientation with cell shape.**

**(A, B)** Distribution of divisions with perimeter-defined cell shape and area-defined cell shape respectively in all cells of the unstretched and stretched control MO, NuMA MO, and rescue tissues. The cell shape axis is mathematically determined based on perimeter ($\theta_P$) and area ($\theta_A$). n=677 cells, 9 embryos; 609 cells, 13 embryos (unstretched, control MO and NuMA MO respectively); 608 cells, 9 embryos; 357 cells, 10 embryos (stretched, control MO and NuMA MO respectively); 304 cells, 11 embryos (rescue). Solid and dashed lines represent median and quartiles respectively.

**(C)** Mean cell area in unstretched and stretched control MO and NuMA MO tissues. n=9 embryos; 13 embryos (unstretched, control MO and NuMA MO respectively); 9 embryos; 10 embryos (stretched, control MO and NuMA MO respectively). Error bars represent mean ± SEM. Data in A, B, and C were analysed using Kruskal-Wallis test and Dunn's multiple comparisons test.

**(D)** Distribution of division orientation with TCV-defined cell shape against cell area in unstretched and stretched control MO and NuMA MO tissues. Data were analysed using Spearman rank correlation test. n=677 cells, 9 embryos; 609 cells, 13 embryos (unstretched, control MO and NuMA MO respectively); 608 cells, 9 embryos; 357 cells, 10 embryos (stretched, control MO and NuMA MO respectively). Statistical significance: $p<0.001$***, $p<0.01$**, $p<0.05$*.

### Videos

**Video 1. Localisation of NuMA in the animal cap.** Time-lapse confocal microscopy of cells in the superficial layer of an unstretched *Xenopus laevis* animal cap explant. The cells are expressing GFP-NuMA (cyan), chromatin marker (mCherry-histone2B; magenta), and a membrane marker (BFP-CAAX; yellow). The white box marks the cell shown in Fig. 1A and Video 2. Time is shown as hour: minute: second. Time interval, 30 s; video frame rate, 20 fps; scale bar, 20 µm.

**Video 2. Localisation of NuMA in a mitotic cell.** Time-lapse confocal microscopy of a cell at interphase and mitosis in an unstretched *Xenopus laevis* animal cap explant. The cell is expressing GFP-NuMA (cyan/inverted grayscale), chromatin marker (mCherry-histone2B; magenta), and a membrane marker (BFP-CAAX; yellow). Time is shown as minute: second. Time interval, 30 s; video frame rate, 10 fps; scale bar, 10 µm. Related to Fig. 1A.

**Video 3. Spindle oscillation.** Time-lapse confocal microscopy of a cell undergoing spindle oscillations in a stretched *Xenopus laevis* animal cap explant. The cell is expressing a membrane marker (GFP-CAAX; green), chromatin marker (mCherry-histone2B; magenta), and contains control morpholino. White arrow marks the duration of the oscillation. Time is shown as minute: second. Time interval, 5 s; video frame rate, 20 fps; scale bar, 10 µm. Related to Fig. 2A and Fig. S2G.

**Video 4. Non-oscillating spindle.** Time-lapse confocal microscopy of a cell with a non-oscillating spindle in a stretched *Xenopus laevis* animal cap explant. The cell is expressing a membrane marker (GFP-CAAX; green), chromatin marker (mCherry-histone2B; magenta), and contains control morpholino. Time is shown as minute: second. Time interval, 5 s; video frame rate, 20 fps; scale bar 10 µm.

**Video 5. Division orientation in the animal cap tissue.** Time-lapse confocal microscopy of cells in the superficial layer of a stretched *Xenopus laevis* animal cap explant. The cells are expressing a microtubule marker (GFP-α-tubulin; green), chromatin marker (mCherry-histone2B; magenta), and contain control morpholino. Upper white box indicates the cell in Fig. 3B and Video 6. Lower white box indicates the cell in Fig. 3B' and Video 7. Time is shown as hour: minute: second. Time interval, 30 s; video frame rate, 20 fps; scale bar 50 µm.

**Video 6. Division orientation parallel to the stretch axis.** Time-lapse confocal microscopy of a cell dividing along the axis of stretch in the *Xenopus laevis* animal cap explant. The cell is expressing a microtubule marker (GFP-α-tubulin; green), chromatin marker (mCherry-histone2B; magenta), and contains control morpholino. Time is shown as minute: second. Time interval, 30 s; video frame rate, 7 fps; scale bar 10 µm. Related to Fig. 3B.

**Video 7. Division orientation perpendicular to the stretch axis.** Time-lapse confocal microscopy of a cell dividing perpendicular to the axis of stretch in the *Xenopus laevis* animal cap explant. The cell is expressing a microtubule marker (GFP-α-tubulin; green), chromatin marker (mCherry-histone2B; magenta), and contains control morpholino. Time is shown as minute: second. Time interval, 30 s; video frame rate, 7 fps; scale bar 10 µm. Related to Fig. 3B'.
